# Feedback inhibition of Ras activity coordinates cell fusion with cell-cell contact

**DOI:** 10.1101/152041

**Authors:** Laura Merlini, Bita Khalili, Omaya Dudin, Laetitia Michon, Vincent Vincenzetti, Sophie G Martin

## Abstract

In fission yeast *Schizosaccharomyces pombe*, pheromone signalling engages a GPCR-Ras-MAPK cascade to trigger sexual differentiation leading to gamete fusion. Cell-cell fusion necessitates local cell wall digestion, the location of which relies on an initially dynamic actin fusion focus that becomes stabilized upon local enrichment of the signalling cascade. We constructed a live-reporter of active Ras1 (Ras1-GTP), also functional in *S. cerevisiae*, which revealed Ras activity at polarity sites peaking on the fusion structure before fusion. Remarkably, constitutive Ras1 activation promoted fusion focus stabilization and fusion attempts irrespective of cell-cell pairing, leading to cell lysis. Ras1 activity is restricted by the GTPase activating protein (GAP) Gap1, itself recruited to sites of Ras1-GTP. While the GAP domain on its own does not suffice for this localization, its recruitment to Ras1-GTP sites is essential to block untimely fusion attempts. We conclude that negative feedback control of Ras activity restrains the MAPK signal and couples fusion with cell-cell engagement.

## Introduction

Feedback inhibition is a very common mechanism to regulate signaling pathways, with various possible outcomes. Negative feedbacks drive the oscillatory behaviors of the cell cycle, circadian clocks, or synthetic networks (Elowitz and Leibler, 2000; Ferrell, 2013; Novak and Tyson, 2008). Negative feedbacks also play a fundamental role in conferring robustness to signal transduction cascades, by dynamically adjusting the system’s output. One example of this is the coordinated synthesis of all ribosomal components (Nomura et al., 1984). These principles are used for temporal control of signaling pathways, but also to produce spatial patterns both during multicellular development (Ribes and Briscoe, 2009) and in single cells (Wu and Lew, 2013). During cell polarization, negative feedbacks have been shown to counteract the positive feedbacks that promote the formation of a single site of polarization, thus providing dynamic adaptation (Das et al., 2012; Ozbudak et al., 2005), limiting the size of the growth zone (Hwang et al., 2008), promoting robustness against variation in polarity factor concentration (Howell et al., 2012), or promoting a morphogenetic transition (Okada et al., 2013). Here, we describe a spatial negative feedback that defines the timing of cell-cell fusion, by coordinating the achievement of a stable polarization state with that of the partner cell.

We are interested in understanding the cellular events driving cell-cell fusion, a process that strongly relies on positive feedback. We have recently described the course of events that leads two haploid fission yeast (*Schizosaccharomyces pombe*) cells of opposite mating type to form a diploid zygote. Early during the mating process, the haploid partners exhibit a polarity patch of active Cdc42 GTPase that dynamically forms and disassembles, exploring multiple sites at the cell cortex (Bendezu and Martin, 2013). This dynamic patch is a site of both own-pheromone secretion and sensing of the opposite-type pheromone (Merlini et al., 2016). Because patch lifetime is prolonged upon higher pheromone perception, adjacent patches in cells of opposite mating type positively feedback to stabilize each other. This leads to cell pairing and growth of the two haploid cells towards their partner.

This positive feedback is maintained and amplified during the fusion process, which requires a dedicated actin-based aster – the fusion focus – nucleated by the formin Fus1 (Dudin et al., 2015; Petersen et al., 1995). The fusion focus underlies the concentration of the pheromone secretion and perception machineries at a focal point. Reciprocally, local activation of the downstream MAPK signaling cascade spatially constrains the focus, leading to its immobilization at facing locations in the two partner cells, now committed to fusion (Dudin et al., 2016). Focus immobilization drives fusion because it allows the type V myosin motor Myo52 to deliver cell wall digestive enzymes at a precise location to locally pierce through the cell wall and permit plasma membrane merging (Dudin et al., 2015; Dudin et al., 2017). Fusion commitment, and thus cell wall digestion, does not directly require cell-cell contact. Indeed, forcing the positive feedback to occur in a single cell, by engineering autocrine cells that respond to self-produced pheromones, leads to fusion attempts without a partner. This causes cell lysis because the locally digested cell wall no longer resists the strong internal turgor pressure (Dudin et al., 2016). Yet, lysis is extremely rare during wildtype cell fusion, suggesting the existence of mechanisms to couple fusion commitment with the formation of cell pairs.

A Ras-MAPK signaling pathway, functionally homologous to the mammalian Ras-Raf-MEK-ERK mitogenic pathway (Hughes et al., 1993), is intimately involved in controlling fission yeast mating. The single *S. pombe* Ras protein, Ras1, whose activated form directly binds the MAP3K Byr2 (Masuda et al., 1995), is an essential activator of the MAP kinase cascade that transduces the pheromone signal (Fukui et al., 1986; Wang et al., 1991b). Two Guanine nucleotide Exchange Factors (GEFs) promote Ras1 activation: a constitutively expressed GEF Efc25, which activates Ras1 for cell polarization during mitotic growth (Papadaki et al., 2002), and a pheromone-induced GEF Ste6, which is required for Ras1 activation of the MAPK cascade during mating (Hughes et al., 1994). Dependence of Ste6 transcription on the MAPK signal creates an additional positive feedback (Hughes et al., 1994; Mata and Bahler, 2006). A single GTPase Activating Protein (GAP), Gap1, is predicted to promote GTP hydrolysis and return Ras1 to its inactive state (Imai et al., 1991; Wang et al., 1991a; Weston et al., 2013). Interestingly, hyper-activation of Ras1, like its deletion, causes sterility, but with distinct phenotype: while *ras1Δ* cells do not engage in mating (Fukui et al., 1986), constitutively active alleles of Ras1 and deletion of *gap1* provoke cell death during mating, which was recently proposed to result from unsustainable cell elongation from multiple sites (Weston et al., 2013). We made the alternative hypothesis that this phenotype is caused by premature fusion attempts. Below, we show that the Ras GAP Gap1 forms a negative feedback that restricts Ras1 activation to sites of pheromone signaling, drives dynamic polarization and prevents fusion commitment during early mating stages to couple it with cell-cell pairing.

## Results

### Constitutive Ras activation promotes untimely fusion attempts

Inactivation and hyper-activation of Ras1, the sole Ras GTPase in fission yeast, both lead to sterility, but exhibit distinct phenotypes (Fukui et al., 1986; Nadin-Davis et al., 1986): Whereas *ras1*Δ remained as short unpolarized cells when placed in mating conditions, homothallic cells carrying a GTP-locked Ras1 allele (*ras1^G17V^* or *ras1^Q66L^*) extended mating projections at apparent aberrant locations and often lysed (Figure 1A) (Nadin-Davis et al., 1986; Weston et al., 2013). Similarly, *h- ras1^G17V^* or *ras1^Q66L^* cells lacking the P-factor protease Sxa2 readily extended mating projections and lysed upon synthetic P-factor exposure, whereas wildtype cells did not lyse, as shown previously (Figure 1B, MovieS1) (Dudin et al., 2016; Weston et al., 2013). Importantly, cell lysis was suppressed by deleting *fus1* in these cells, suggesting that lysis may arise from an untimely fusion attempt (Figure 1B).

**Figure 1:**
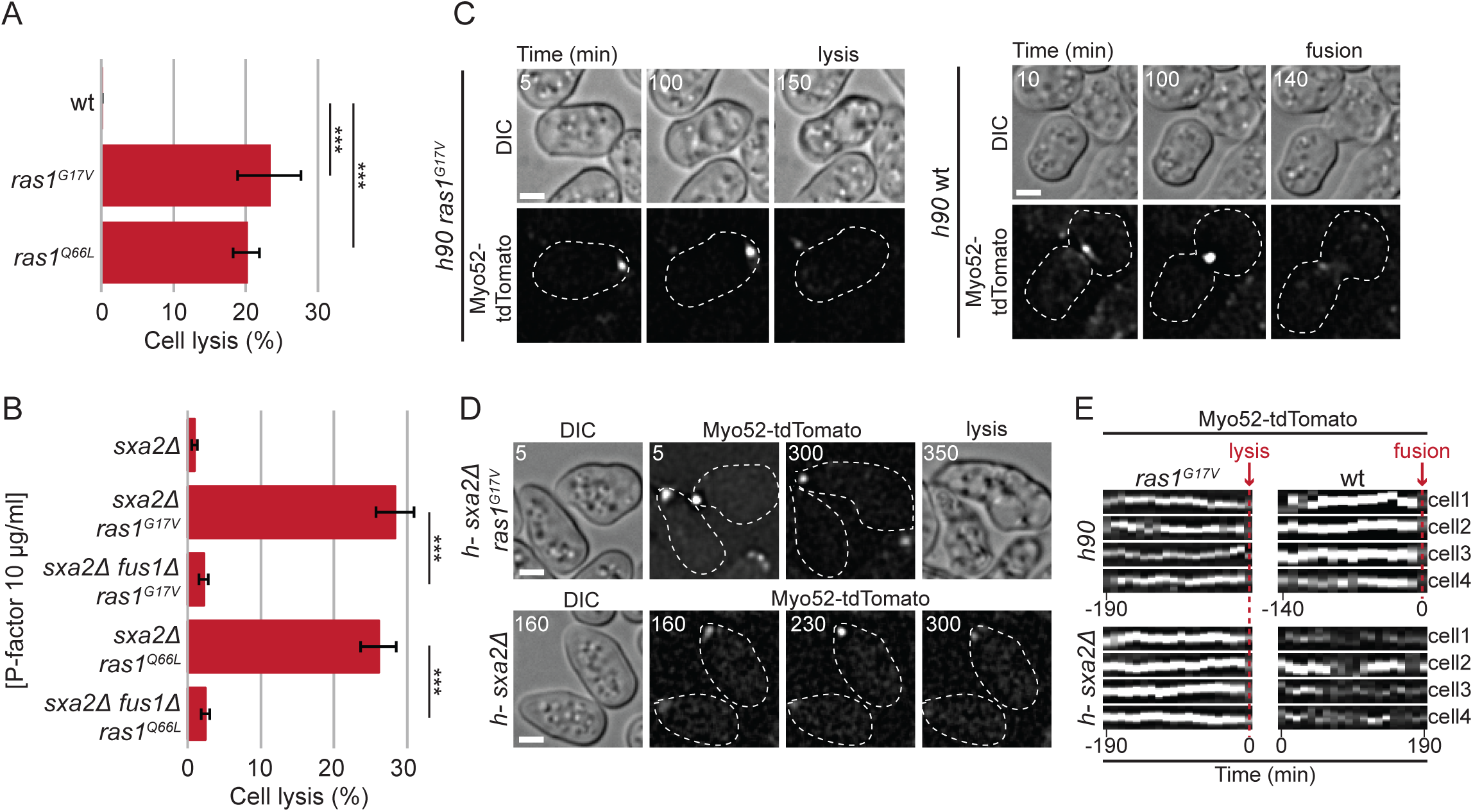
Constitutive Ras activation promotes untimely fusion attempts. (A) Percentage of cell lysis of homothallic wt and constitutively active *ras* mutants after 14h in MSL-N (n > 500 for 3 independent experiments); *** indicates 5.85 x 10^-6^ ≤ p-value^t-test^ ≤ 1.1 x 10^-5^. (B) Percentage of cell lysis of heterothallic *h- sxa2Δ*, *h- sxa2Δ ras1^G17V^* and *h- sxa2Δ ras1^Q66L^* cells, with or without *fus1* deletion (*fus1Δ*), 14h after 10μg/ml synthetic P-factor addition in MSL-N (n > 500 for 3 independent experiments); *** indicates 4.58 x 10^-6^ ≤ p-value^t-test^ ≤ 1.43 x 10^-5^. (C) DIC and Myo52-tdTomato time-lapse images of homothallic *ras1^G17V^* and wt during mating. Note the persistent Myo52 focus until cell lysis in the unpaired *ras1^G17V^* cell, whereas the Myo52 focus is only observed in cell pairs during fusion in wt cells. The time of cell lysis (*ras1^G17V^*) and fusion (wt) are indicated. (D) DIC and Myo52-tdTomato time-lapse images of heterothallic *h- sxa2Δ ras1^G17V^* and *h- sxa2Δ* cells treated with 10μg/ml P-factor. Note the persistent Myo52 focus and cell lysis in *ras1^G17V^* cells, and the unstable Myo52 signal in wt cells. (E) Kymographs of four representative cell tips showing the formation of a stable Myo52 focus in *h90 ras1^G17V^* mating cells and *h- sxa2Δ ras1^G17V^* mutant cells exposed to 10μg/ml P-factor. The kymographs are aligned to the time of lysis. By contrast, *ras1^+^* cells form a focus late in the fusion process (in *h90*, the kymographs are aligned to the time of fusion) or only transiently (in *h- sxa2Δ* exposed to P-factor; no alignment of the kymographs). Bars = 2 μm. Time is in minutes from the start of imaging.

Consistent with this hypothesis, cells with constitutive Ras1 activation displayed a strong, focal fluorescent signal of Myo52-tdTomato, reminiscent of that observed at the fusion focus of wildtype cell pairs (Dudin et al., 2015). This signal formed and remained stable over long time periods in unpaired homothallic cells prior to cell lysis (Figure 1C, E). By contrast, wildtype cells form a fusion focus only after pairing (Figure 1C, E). Similarly, in heterothallic *sxa2*Δ cells exposed to synthetic pheromone, a stable Myo52 focus was formed upon constitutive Ras1 activation, whereas the Myo52 signal was broad and only transiently focalized in *ras1^+^* cells (Figure 1D-E, Movie S1). These observations suggest Ras1 activation promotes fusion focus stabilization for cell fusion.

### RasAct – a probe for in situ labeling of active GTP-bound Ras

To define the cellular location of Ras activity, we developed a fluorescent probe that would specifically label the GTP-bound form of Ras1. Ras1-GTP directly associates with the MAP3K Byr2 (Masuda et al., 1995), and the structure of the Byr2 Ras-GTP binding domain (RBD) has been solved (Gronwald et al., 2001). We cloned three tandem repeats of the Byr2 RBD followed by three tandem repeats of GFP and expressed this probe, called RasAct^GFP^ below, under control of the constitutive *pak1* promoter (Figure 2A). In vegetative cells, RasAct^GFP^ localized to cell poles and septa, where Ras1 is predicted to be active. Pole and septum localizations were abolished in cells carrying a deletion of *ras1* or *efc25*, encoding the sole Ras GEF expressed during vegetative life stages (Papadaki et al., 2002), or a GDP-locked *ras1^S22N^* allele (Figure 2B, D). RasAct^GFP^ also accumulated in the nucleus, though this localization was not affected by Ras1 activity state, suggesting this is a spurious localization (Figure 2B). In contrast to the pole-restricted RasAct^GFP^ localization, GFP-Ras1 decorated the entire cell cortex and was only weakly enriched at cell poles (Figure 2C). We conclude that cortical RasAct^GFP^ localization reports on sites of Ras1-GTP, which represents only a fraction of total Ras1.

**Figure 2:**
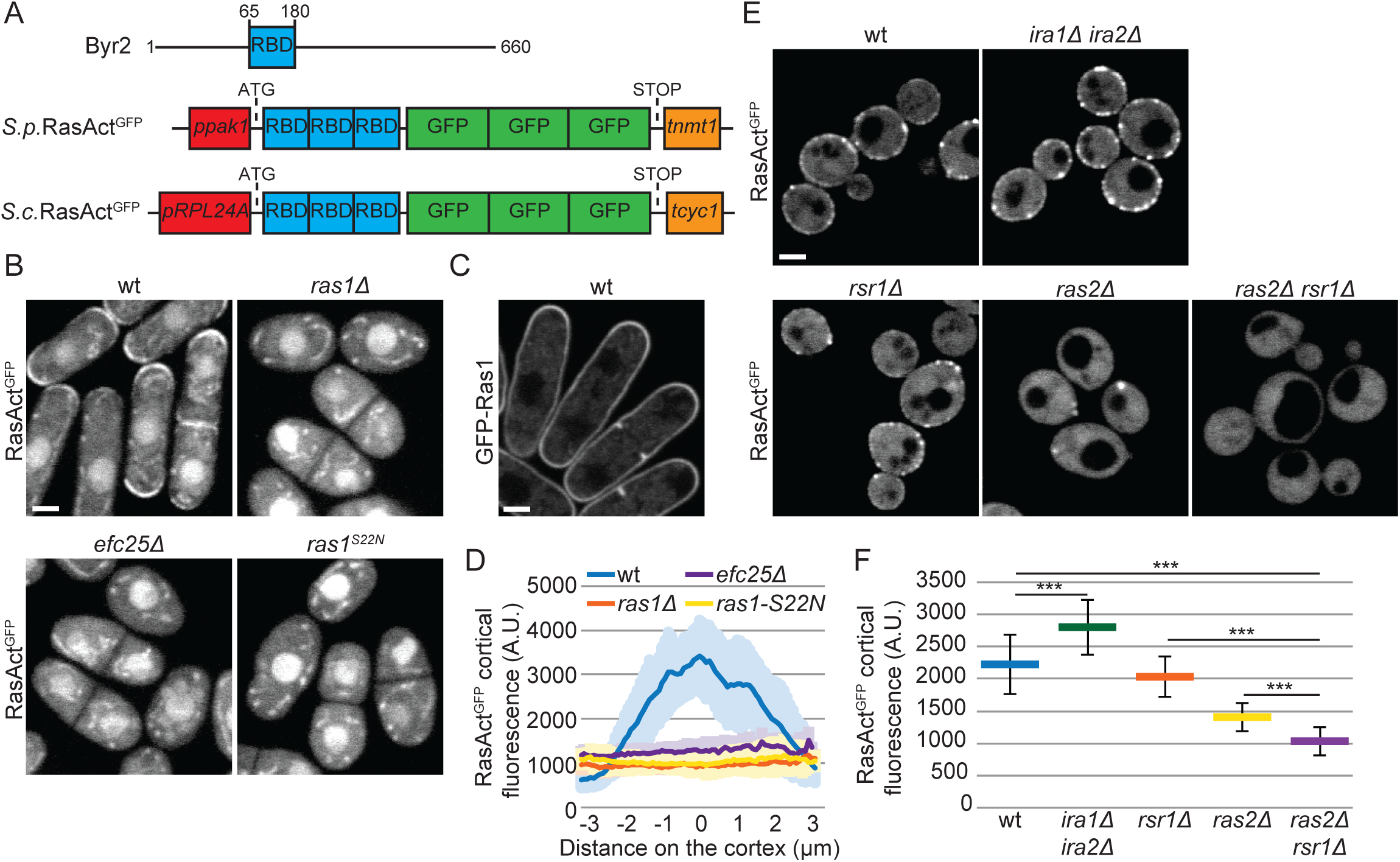
A probe for the visualization of Ras-GTP. (A) Schematic representation of the MAP2K protein Byr2 (top) and of the probes for the visualization of Ras-GTP (RasAct^GFP^) in *Schizosaccharomyces pombe* (*S. p.* middle) and *Saccharomyces cerevisiae* (*S. c.* bottom). RBD = Ras-GTP Binding Domain, GFP = Green Fluorescent Protein. Promoters and terminators used for gene expression are indicated. (B, C) Localization of RasAct^GFP^ (B) and GFP-Ras1 (C) during vegetative growth of fission yeast cells of indicated genotypes. (D) Cortical profiles of RasAct^GFP^ fluorescence intensity at the cell tips in strains as in (B); n = 25. (E) Localization of RasAct^GFP^ during vegetative growth of budding yeast cells of indicated genotypes. (F) Average total RasAct^GFP^ cortical fluorescence in strains as in (E); n = 25; *** indicates 3.9 x 10^-13^ ≤ p-value^t-test^ ≤ 3.7 x 10^-5^. Bars = 2 μm.

We tested whether RasAct^GFP^ is able to detect Ras-GTP in other organisms beyond fission yeast. RasAct^GFP^ was expressed in *Saccharomyces cerevisiae* under the control of *pRPL24A* promoter and integrated as single genomic copy (Figure 2A). In budding yeast cells, RasAct^GFP^ localized in the cytoplasm and decorated regions of the cell cortex (Figure 2E). *S. cerevisiae* encodes two Ras isoforms, named Ras1 and Ras2 (Kataoka et al., 1984), both implicated in glucose sensing (Conrad et al., 2014), negatively regulated by two GAPs Ira1 and Ira2 (Tanaka et al., 1990), and localized to the plasma membrane (Manandhar et al., 2010). These cells also express a third Ras-like protein, Rsr1, involved in cell polarity regulation (Bi and Park, 2012). Interestingly, deletion of *ira1* and *ira2* increased RasAct^GFP^ cortical levels (Figure 2E-F), suggesting RasAct^GFP^ reports on Ras-GTP levels. In cells lacking Ras2, RasAct^GFP^ was poorly recruited to the cell cortex (Figure 2E-F). The residual cortical localization was dependent on Rsr1, as RasAct^GFP^ did not label the cell cortex in double *ras2Δ rsr1Δ* mutants. We conclude that the RasAct^GFP^ probe reports on both Rsr1- and Ras2-GTP (Figure 2E-F). These results establish RasAct^GFP^ as a tool to detect local levels of active Ras in both fission and budding yeast cells.

### Ras activity at the fusion site peaks before cell fusion

We used RasAct^GFP^ to probe where Ras1 is active in mating fission yeast cells. During dynamic polarization, RasAct^GFP^ localized to dynamic sites at the cortex, which overlapped with sites of Cdc42 activity, as labeled by the scaffold protein Scd2 (Figure 3A, S1). We have recently shown that GFP-Ras1 weakly accumulates at these sites, but is also present broadly at the cell cortex (Merlini et al., 2016). Timelapse imaging showed simultaneous accumulation and loss of RasAct^GFP^ and Scd2-mCherry signals at dynamic cortical sites, in agreement with Ras1 acting as activator of Cdc42 signaling (Chang et al., 1994; Merlini et al., 2016). Thus, Ras1-GTP, like Cdc42-GTP, exhibits oscillatory polarization dynamics during early mating.

**Figure 3:**
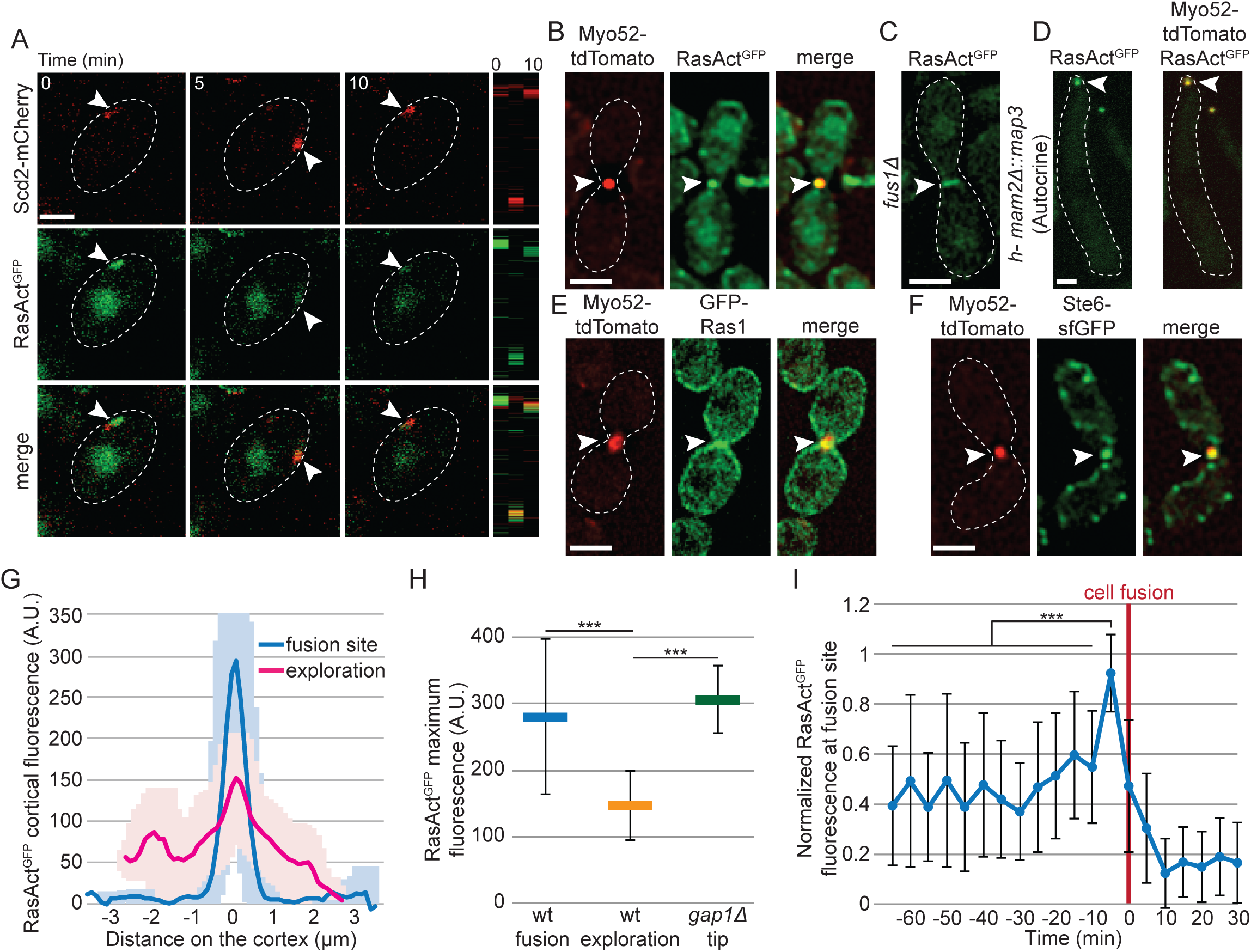
Ras1 is active at polarity sites and at the fusion focus. (A) Spinning disk confocal projections of *h90* wild-type strains showing co-localization of Scd2-mCherry and RasAct^GFP^ during early stages of mating. Kymographs of the cell periphery are shown on the right. Arrowheads highlight dynamic zones of co-localization. (B) Co-localization of RasAct^GFP^ (green) with Myo52-tdTomato (red) in homothallic wild type cells. (C) Localization of RasAct^GFP^ in *fus1*Δ cell pair. (D) Co-localization of RasAct^GFP^ (green) with Myo52-tdTomato (red) in *mam2Δ::map3* autocrine M-cells. (E) Co-localization of GFP-Ras1 (green) with Myo52-tdTomato (red) in homothallic wild type strains. (F) Co-localization of Ste6-sfGFP (green) with Myo52-tdTomato (red) in homothallic wild type cells. In B-F, arrowheads highlight zones of localization and co-localization with Myo52-tdTomato. (G) Cortical profiles of RasAct^GFP^ fluorescence in homothallic wt strains during early (exploration) or late (fusion site) mating stages, showing that Ras activity is enriched at the time of fusion; n = 20. The thick line is the average and the corresponding shaded area the standard deviation. (H) Quantification of maximal RasAct^GFP^ cortical fluorescence in homothallic wt cells during early (exploration) or late (fusion) mating stages and at the mating projections of homothallic *gap1Δ* cells. Average values of the 5 brightest pixels are shown; n = 20; *** indicates 9.7 x 10^-34^ ≤ p-value^t-test^ ≤ 2.8 x 10^-13^. (I) Average normalized value of RasAct^GFP^ cortical fluorescence over time at the fusion site in homothallic wt cells. All fluorescence profiles were aligned to the fusion time (t=0), as assessed by diffusion of a cytosolic marker from one cell to the other (Vjestica et al., 2016); n = 22. Each profile was normalized to its maximal value. *t* test is calculated between each of the first 12 timepoints and the time before fusion (-5 min), *** indicates 8.7 x 10^-12^ ≤ p-value^t-test^ ≥ 1.1 x 10^-5^ after Bonferroni correction. Bars = 2 μm.

During fusion, RasAct^GFP^ strongly accumulated at the fusion focus: RasAct^GFP^ signal co-localized with Myo52 and its concentration at a focal site depended on Fus1 (Figure 3B-C). RasAct^GFP^ was also concentrated at the fusion focus in autocrine M-cells that attempt cell fusion in absence of a partner cell (Figure 3D). By contrast, GFP-Ras1 was enriched at the mating projection over a broader zone (Figure 3E), suggesting that Ras1-GTP is restricted to the fusion site with Ras1-GDP present in surrounding regions. During mating, activation of Ras1 is thought to occur downstream of pheromone receptor signaling, which takes place locally first at dynamic polarization sites (Merlini et al., 2016) and then at the fusion focus (Dudin et al., 2016). Unfortunately, tagging of the pheromone-induced Ras GEF Ste6 (Hughes et al., 1990; Papadaki et al., 2002) with GFP or sfGFP at the N- or C-terminus produced a protein with reduced function, as these cells exhibited only 25% mating efficiency (compared to 75% for wildtype cells in our assay conditions), and we were unable to detect it during dynamic polarization. However, partly functional Ste6-sfGFP colocalized with the fusion focus (Figure 3F), consistent with local activation of Ras1 at this location. We note that Efc25-GFP was undetectable during mating (data not shown). We conclude that Ras1 is locally activated first at dynamic polarization sites and then at the fusion site.

The spatial distribution of cortical RasAct^GFP^ was distinct at exploratory zones and the fusion focus: RasAct^GFP^ formed low-intensity, broad peaks at exploratory zones; it formed higher-intensity, sharper ones at the fusion site (Figure 3G-H). Timelapse acquisition before fusion and data alignment to the fusion time further revealed that RasAct^GFP^ levels were highest at the fusion focus immediately before the fusion event (Figure 3I). Thus, local Ras1-GTP concentration peaks at the fusion site just prior to fusion.

### Gap1 GTPase Activating Protein restricts Ras1 activity

Because Ras1 activation peaks before fusion and constitutive Ras1 activation promotes untimely fusion attempts, we asked how Ras activity is controlled to induce fusion. Ras1-GTP hydrolysis is likely to be catalyzed by the GAP Gap1 (Imai et al., 1991; Wang et al., 1991a; Weston et al., 2013). Indeed, we found that MBP-Gap1 accelerated GTP hydrolysis on Ras1 *in vitro*, whereas MBP-Gap1^R340A^ carrying a point mutation in the GAP domain predicted to affect GTP hydrolysis (Sermon et al., 1998) had no effect (Figure 4A). Recombinant ^GST^RBD (a derivative of RasAct^GFP^, expressing just one copy of the Ras-GTP Binding Domain fused to GST; (Kae et al., 2004)) pulled down vast excess of Ras1-GTP from *gap1Δ* extracts, as compared to wildtype extracts, indicating that Gap1 promotes Ras1-GTP hydrolysis also *in vivo* (Figure 4B). In cells lacking Gap1, RasAct^GFP^, like GFP-Ras1, decorated the entire cortex of vegetative growing cells and was also absent from the nucleus (Figure 4C). Thus, most, if not all, Ras1 is in the active state and recruits all RasAct^GFP^ to the plasma membrane. In *gap1Δ* cells during mating, we did not observe discrete dynamic RasAct^GFP^ zones. Instead, RasAct^GFP^ was distributed homogeneously at the cortex in unpolarized cells. In those extending a growth projection, RasAct^GFP^ was broadly distributed around the projection tip, but largely excluded from the back of the cell. This localization mimics that of GFP-Ras1, again suggesting that most, if not all, Ras1 molecules are active in this mutant (Figure 4C). Fluorescence intensity measurements further showed that RasAct^GFP^ levels at the projection tip of *gap1Δ* cells, were significantly higher than those observed during early mating in wildtype cells, and similar to those observed at the fusion focus of wildtype cells engaged in fusion (Figure 3H). Thus, Gap1 is a Ras1 GAP and restricts Ras1 activity.

**Figure 4:**
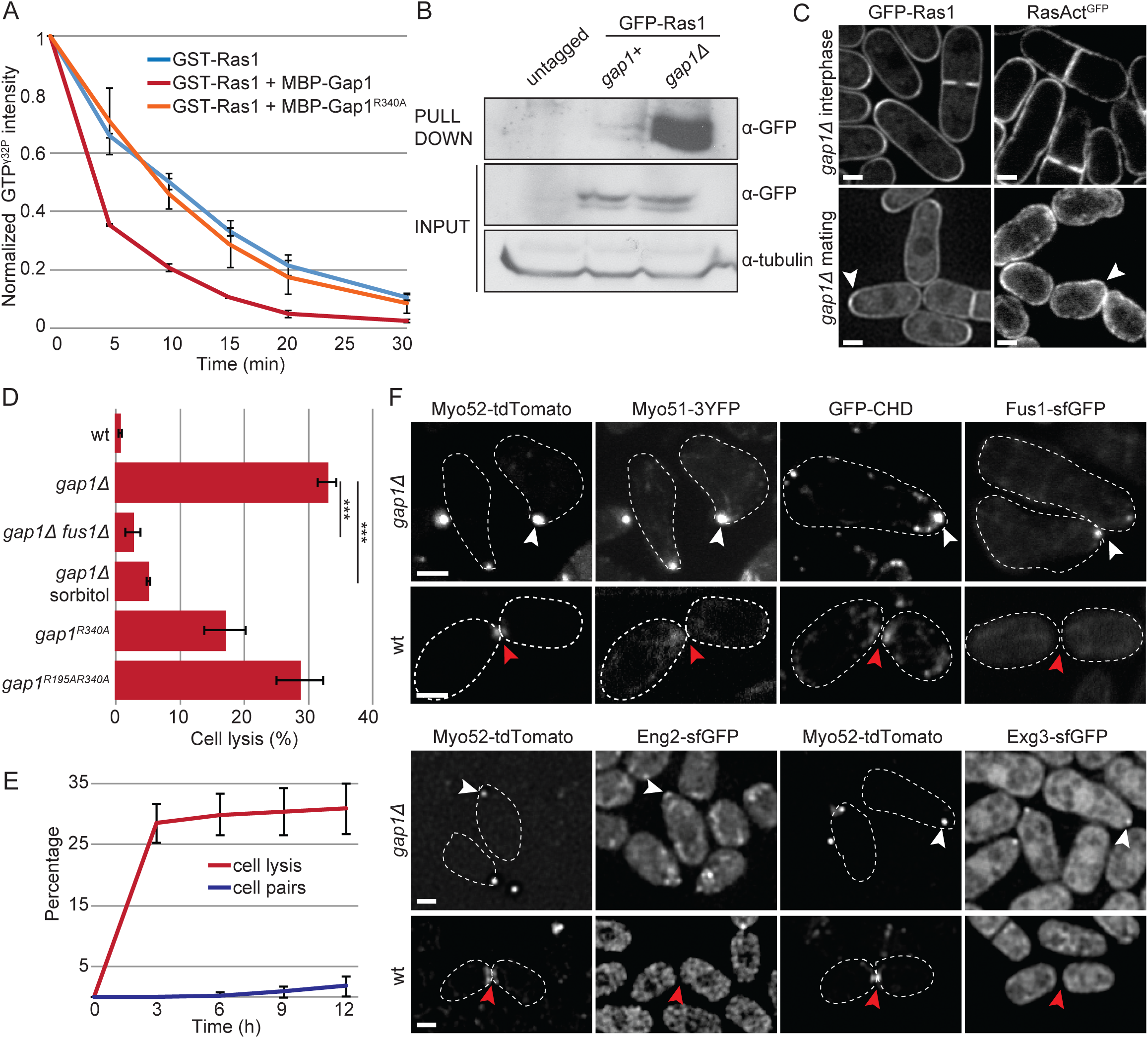
Gap1 is a GTPase Activating Protein for Ras1. (A) *In vitro* GAP assays show that the rate of GTP hydrolysis of purified GST-Ras1-GTP^γ32P^ is increased in the presence of purified MBP-Gap1, but not catalytically inactive MBP-Gap1^R340A^; n = 3. (B) GST-RBD pull-down of protein extracts from *h-* vegetative growing cells of indicated genotypes showing higher GFP-Ras1-GTP levels in *gap1Δ* cells. (C) Localization of GFP-Ras1 (left) and RasAct^GFP^ (right) in *gap1*Δ strains during vegetative growth (top) and mating (bottom). Arrowheads indicate the shmoo tip. (D) Percentage of cell lysis of homothallic wt and *gap1* mutant (*gap1^R340A^, gap1^R195A R340A^* and *gap1Δ*) cells treated with 1.2M sorbitol or lacking *fus1* (*fus1Δ*) after 14h in MSL-N (n > 500 for 3 independent experiments); *** indicates 9.34 x 10^-7^ ≤ p-value^t-test^ ≤ 6.54 x 10^-6^. (E) Percentage of cell lysis and cell pair formation in homothallic *gap1Δ* cells placed on MSL-N over time (n > 200 for 3 independent experiments). (F) Type V Myosins (Myo52-tdTomato and Myo51-3YFP), actin (GFP-CHD), formin Fus1-sfGFP and cell wall glucanases (Eng2-sfGFP and Exg3-sfGFP) are focalized at the shmoo tip of unpaired *gap1Δ* cells (white arrowheads) but either not detectable or not focalized (red arrowheads) at the shmoo tips of wt cells not yet engaged in fusion. Wildtype cells focalize these factors at later stages during the fusion process (not shown; (Dudin et al., 2015)). Bars = 2 μm. Plots show averages and standard deviations.

Multiple lines of evidence showed that cells lacking Gap1 or its GAP activity, like *ras1^Q66L^* cells, engaged into untimely fusion attempts. We have recently described that *gap1Δ* cells display diminished dynamic polarization of Cdc42 and stabilize a single site of growth at reduced pheromone concentrations, leading to misoriented growth projections (Merlini et al., 2016), thus reducing cell-cell pairing efficiency (Figure 4E). Remarkably, homothallic cells with a *gap1* deletion or point mutation abolishing the GAP activity (*gap1^R340A^, gap1^R195A R340A^*) often lysed (Weston et al., 2013) (Figure 4D). We note that cell lysis was slightly less frequent in cells carrying the mutation *gap1^R340A^*, predicted to block GTP hydrolysis, than *gap1^R195A R340A^*, predicted to also block Ras1-GTP binding (Scheffzek et al., 1997; Sermon et al., 1998), suggesting that both Ras1-GTP binding and hydrolysis contribute to Gap1 function *in vivo*; notably, 34.6 ± 4.1% of *gap1^R340A^* were also able to mate. Lysis of cells lacking *gap1* was an early event, with almost maximal levels reached already 3h after mating induction (Figure 4E). Importantly, osmo-stabilization by addition of 1.2M sorbitol efficiently prevented lysis (Figure 4D, S2A), indicating lysis was due to cell wall degradation. Furthermore, *fus1* deletion suppressed lysis, though it did not prevent the occurrence of misoriented growth projections (Figure 4D, S2A). This suggests that lysis is due to precocious assembly of the fusion focus.

In wildtype cells, the formation of the actin fusion focus takes place only after cells are paired, with single cells or even cells that have just paired not yet exhibiting a tight concentration of type V myosins (Figure 4F) (Dudin et al., 2015). Strikingly, cells lacking Gap1 assembled at the tip of unpaired cell projections a focal structure labeled by markers of the fusion focus: the formin Fus1, GFP-CHD-labeled F-actin, the type V myosins Myo52 and Myo51, as well as the glucanases Eng2 and Exg3 for cell wall digestion (Figure 4F, Movie S2, S3) (Dudin et al., 2015). In addition, heterothallic *h- sxa2Δ gap1Δ or h+ sxa1Δ gap1Δ* cells exposed to synthetic P- or M-factor, respectively, also readily assembled a stable fusion focus and lysed (Figure S2B-E). Again, lysis was prevented by *fus1* deletion (Figure S2C, E). Finally, *gap1* deletion increased the percentage of lysing autocrine M-cells, in which pheromone signaling focalization occurs through a cell-autonomous positive feedback (Dudin et al., 2016) (Figure S2F). We conclude that, by promoting Ras1-GTP hydrolysis, Gap1 protects cells against untimely fusion attempts.

### Ras1 activity promotes MAPK focalization

We recently showed that focalization of pheromone signaling, including the pheromone receptors, the coupled Gα and the MAPK cascade, at the fusion focus, promotes cell fusion by stabilizing the focus (Dudin et al., 2016). Because *gap1Δ* cells engage in precocious fusion events, we tested the spatial organization of these signaling molecules. Whereas pheromone signaling components accumulate at the fusion focus only in committed cell pairs, late in the fusion process (Figure 5A) (Dudin et al., 2016), the MAP2K Byr1 and the M-factor pheromone receptor Map3, localized to the premature fusion focus in unpaired *gap1Δ* cells (Figure 5A). We conclude that constitutive Ras1 activation engages the positive feedback leading to focalization of the pheromone signaling cascade and stabilization of the fusion focus.

**Figure 5:**
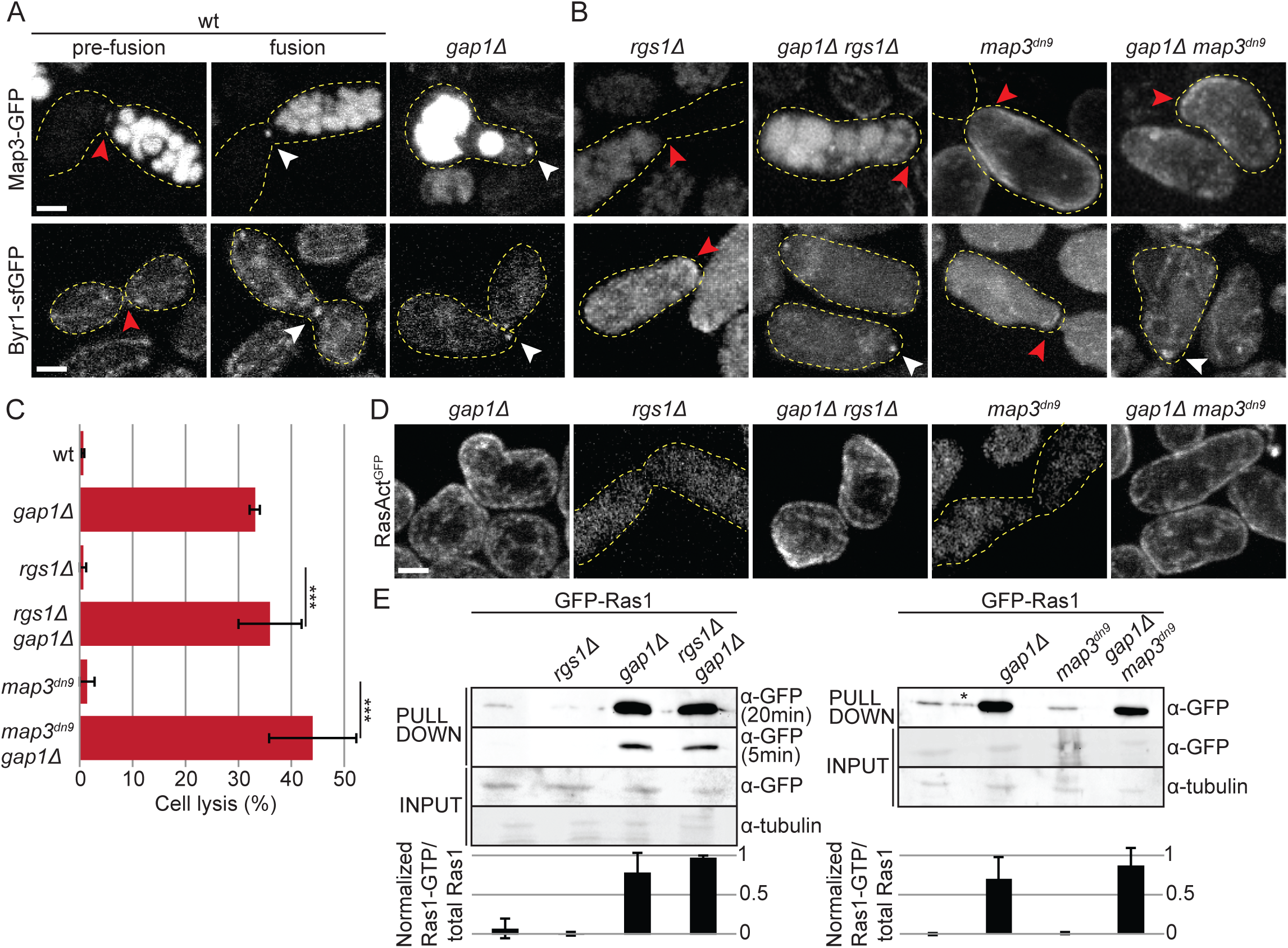
Ras activity promotes MAPK focalization. (A-B) Localization of pheromone receptor Map3-GFP (top) and MAP2K Byr1-sfGFP (bottom) in wt cell pairs not yet engaged in fusion or during the fusion process (left), and at the projection tip of mutant strains of indicated genotypes (right). White arrowheads highlight focalization and red arrowheads zones of broad or undetectable localization. Note that deletion of *gap1* leads to Byr1 MAP2K focalization whether the Map3 receptor is itself focalized or not. (C) Percentage of cell lysis of homothallic wt, *rgs1Δ* and *map3^dn9^* mutants, with or without *gap1* deletion (*gap1Δ*) after 14h in MSL-N (n > 500 for 3 independent experiments). *** indicates p-value^t-test^ ≤ 2.6 x 10^-5^. (D) Localization of RasAct^GFP^ at the mating projections of mutant strains of indicated genotypes. Note undetectable local Ras activation in situations when the pheromone receptor is unfocalized and quasi-uniform cortical activation upon *gap1* deletion. (E) GST-RBD pull-down of protein extracts from *h90* homothallic cells with indicated genotypes after 4h in MSL-N (top). Average intensity quantification from three independent experiments is shown (bottom). * This band, in an empty lane left to space out samples, is due to overflow from neighboring lanes. Bars = 2 μm.

To understand the relationship between Ras1 activation and engagement of the positive feedback, we used two mutant conditions that impair receptor focalization: *map3^dn9^* (a truncation of the receptor C-terminal tail blocking receptor endocytosis; (Hirota et al., 2001)) and *rgs1Δ* (a deletion of the GTPase activating protein of the receptor-associated Gα protein (Pereira and Jones, 2001; Watson et al., 1999)) fail to focalize Map3 receptor and Byr1 MAP2K, and are fusion-defective (Figure 5B) (Dudin et al., 2016). Consistently, in all *rgs1Δ* and in the majority of *map3^dn9^* cells, RasAct^GFP^ did not reveal local Ras activity at the cell-cell contact site (Figure 5D). A small proportion (13 out of 102 cells) of *map3^dn9^* cells localized RasAct^GFP^ over a broad zone at the cell-cell contact (data not shown). However, pull-down assays with ^GST^RBD detected Ras1-GTP at levels similar as in wildtype in these mutants (Figure 5E), suggesting Ras1-GTP is present but distributed over a large area and thus not detected by microscopy. These results are consistent with the view that the narrow distribution of Ras1-GTP follows from the concentration of active receptors at the fusion focus.

Interestingly, deletion of *gap1* in *map3^dn9^* and *rgs1Δ* cells caused constitutive activation of Ras1 on the entire cell cortex, formation of mating projections at aberrant sites and precocious fusion attempts, leading to cell lysis, like *gap1Δ* single mutants (Figure 5C-D). Thus, constitutive Ras1 activation can bypass the normal focalization of the pheromone receptor. Indeed, in these double mutants, the pheromone receptor was still not enriched at the fusion focus but the MAP2K cascade component Byr1 was (Figure 5B), These data indicate that Ras1-GTP promotes MAPK localization to the actin fusion structure independently of the localization of upstream signaling components and does not need to be restricted to the fusion focus to fulfill this function. These data are also consistent with the notion that MAPK focalization is the critical signal for fusion focus stabilization (Dudin et al, 2016).

Altogether, the results above indicate that elevated Ras1 activity levels at the shmoo site promote the local accumulation of the MAPK cascade on the fusion focus for fusion commitment and that the Ras1 GAP Gap1 restricts Ras1-GTP levels until cells are ready for fusion.

### Gap1 locally restricts Ras1 activity through negative feedback

To understand how Gap1 spatially controls Ras1 activity, we examined Gap1 localization. Previous data, using overexpressed constructs, showed that Gap1 is recruited to the cell cortex by Ras1-GTP after pheromone stimulation (Weston et al., 2013). Gap1-GFP tagged as sole copy at the endogenous genomic locus weakly associated with the cell cortex also in vegetative proliferating cells, and was also present in the cytosol. In these cells, Gap1-GFP fluorescence was observed at cell poles in wildtype, but not in *ras1Δ*, *efc25Δ* or *ras1^S22N^* cells, similar to RasAct^GFP^, in agreement with its recruitment to the cell cortex by Ras1-GTP (Figure 6A). During early mating, Gap1-GFP localized to sites of Cdc42 and Ras1 activation, labeled with Scd2-mCherry (Figure 6B). Short-interval timelapse imaging of Gap1 during dynamic polarization revealed that Gap1-GFP was recruited to Scd2 sites and remained associated with the site beyond its disassembly (Figure 6C-D, S3). At the fusion site, Gap1 distribution was significantly broader than that of RasAct^GFP^ (Figure 6E-F). These two observations suggests Gap1 may be recruited by Ras1-GTP, but remain associated with the cell cortex beyond GTP hydrolysis. Thus, Ras1-GTP may recruit its own inhibitor, forming a negative feedback loop. Such negative feedback loop would be in agreement with the role of Gap1 in promoting polarity patch disassembly during dynamic exploration (Merlini et al., 2016). It may also serve to protect against premature fusion attempts and coordinate fusion with cell-cell contact.

**Figure 6:**
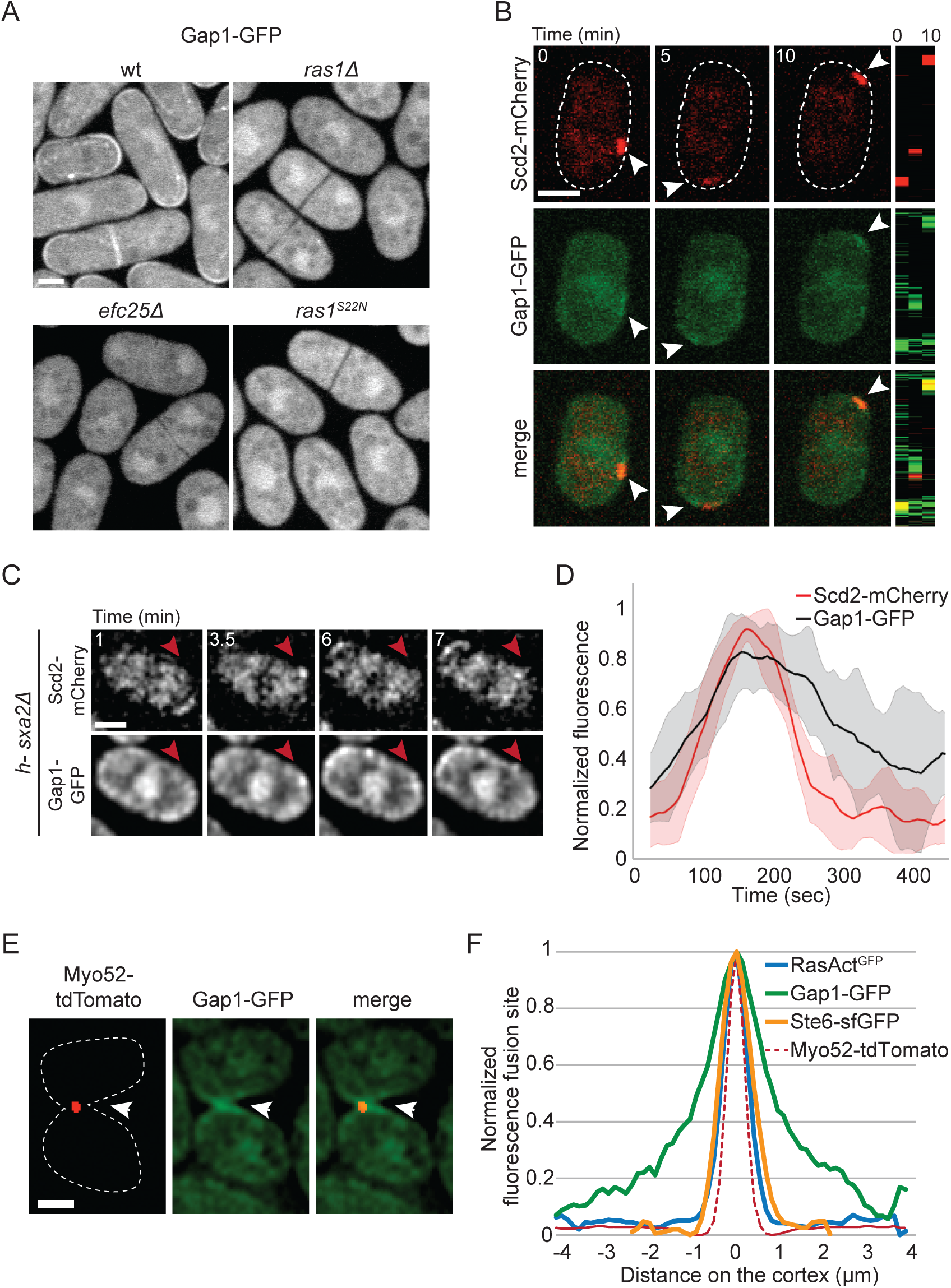
Gap1 is part of a negative feedback on Ras1 activity. (A) Localization of Gap1-GFP during vegetative growth in wt and mutant strains of indicated genotypes. (B) Spinning disk confocal projections of *h90* wild-type strains showing co-localization of Scd2-mCherry (red) and Gap1-GFP (green) during early stages of mating. Kymographs of the cell periphery are shown on the right. Arrowheads highlight dynamic zones of co-localization. (C) Single plane images of *h- sxa2Δ* cells treated with 0.01 μg/ml of synthetic P-factor showing co-localization of Scd2-mCherry and Gap1-GFP in a dynamic polarity patch. The red arrowhead highlights the location of one patch of Scd2-mCherry that appears and disappears in the course of the timelapse and the corresponding location in the Gap1-GFP channel. For more examples, see Figure S3. (D) Cortical profiles of Scd2-mCherry and Gap1-GFP normalized fluorescence intensity at the exploratory patch, showing that Gap1-GFP signal persists after Scd2-mCherry signal has disappeared. Average quantification of 10 patches from 10 different cells is shown. The thick line represents the average and the corresponding shaded area the standard deviation. (E) Spinning disk confocal projections of *h90* wild-type strains expressing Myo52-tdTomato (red) and Gap1-GFP (green). Gap1-GFP signal at the fusion site is broader than the Myo52-tdTomato signal. Arrowheads highlight the localization of Myo52-tdTomato. (F) Cortical profiles of Gap1-GFP, RasAct^GFP^ and Ste6-sfGFP fluorescence at the fusion site in homothallic wt strains as in (E) and Figures 3B and 3F. The profiles were aligned to the Myo52-tdTomato signal co-imaged in the same cells (dashed line); n = 20. Time is in minutes from the start of imaging. Bars = 2 μm.

To test for the existence of, and the possible requirement for, negative feedback, we probed the mode of Gap1 recruitment to Ras1. The recruitment of Gap1 to Ras1-GTP may simply reflect interaction of the GAP domain with Ras1-GTP, as previously suggested (Weston et al., 2013). However, although a wildtype GAP domain was necessary for the cortical localization of Gap1 (Figure S4D), we found that the sole GAP domain (GAP^Gap1^) only weakly localized to the Myo52 focus (Figure 7A), suggesting that N- and C-terminal Gap1 regions also contribute to Gap1 recruitment to sites of Ras1 activity. The GAP^Gap1^ construct, which was expressed under the control of a constitutive promoter in cells lacking the endogenous copy of *gap1* (Figure S4A), did not rescue the lysis phenotype of *gap1Δ* cells (Figure 7B). However, GAP^Gap1^ retained some activity *in vivo*, as it partly rescued cell pairing: about 33% of cells formed pairs and eventually fused 12h after nitrogen starvation. By contrast, full-length Gap1 (Gap1^FL^) expressed under the same conditions and at similar levels (Figure S4A-B), localized broadly at the fusion site and fully rescued the lysis and pairing defects of cells lacking *gap1* (Figure 7A-B). We conclude that robust Gap1 recruitment to Ras1-GTP, necessary to prevent premature fusion attempts, relies on both its GAP domain and additional regions, consistent with negative feedback regulation.

**Figure 7:**
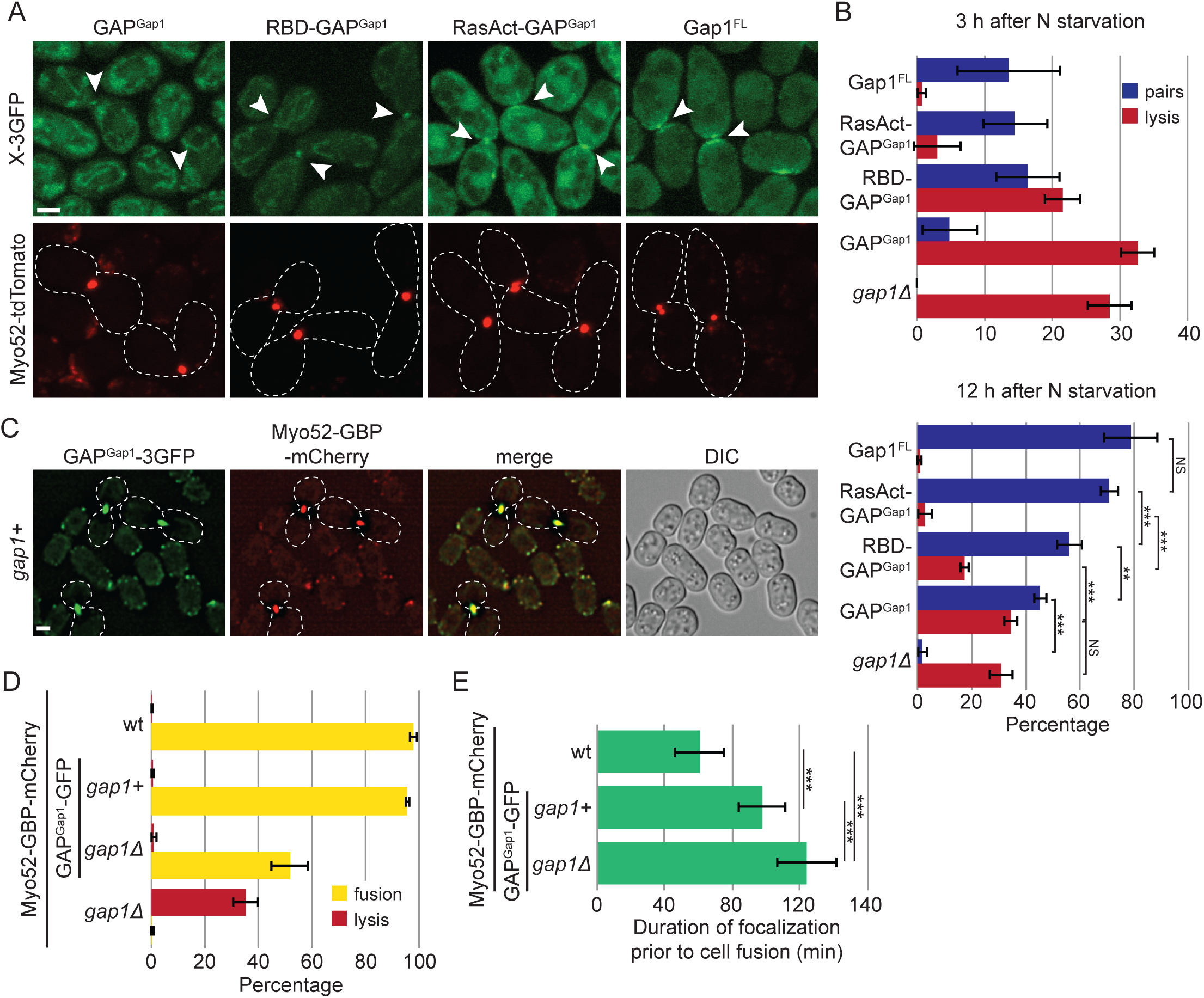
The negative feedback on Ras1 activity couples cell fusion with cell contact. (A) Localization of the sole GAP domain of Gap1 (GAP^Gap1^), this domain fused to a single or triple RBD (RBD-GAP^Gap1^, RasAct-GAP^Gap1^), and full-length Gap1 fused to 3GFP and expressed in *gap1Δ* cells. These constructs are detailed in Fig. S4A. Myo52-tdTomato (red) foci indicate cell pairs in fusion. Arrowheads highlight the localization of the fusion proteins at the Myo52-tdTomato foci. (B) Percentage of cell lysis and cell pair formation of homothallic *gap1Δ* cells expressing or not the fusion constructs shown in (A) 3h (top) and 12h (bottom) after nitrogen starvation (see also Figure 4E; n > 300 for 3 independent experiments). (C) Representative images of GAP^Gap1^-3GFP strains expressing Myo52-GBP-mCherry 8h after nitrogen starvation. GAP^Gap1^-3GFP is recruited to the fusion focus via interaction with Myo52-GBP-mCherry. (D) Percentage of cell lysis and fusion efficiency of wt, *gap1Δ*, GAP^Gap1^-3GFP and *gap1Δ* GAP^Gap1^-3GFP strains expressing Myo52-GBP-mCherry 12h after nitrogen starvation. Note the ability of GAP^Gap1^-3GFP recruitment to the fusion focus to suppress the *gap1Δ* cell lysis. (E) Duration of Myo52-GBP-mCherry focalization before fusion, as assessed by the disassembly of Myo52 focus in strains of indicated genotypes; n = 25. *** indicates 1.2 x 10^-16^ ≤ p-value^t-test^ ≤ 7.2 x 10^-7^. Note the extended fusion time upon recruitment of GAP^Gap1^-3GFP to the fusion focus. Bars = 2 μm.

We engineered recruitment of the GAP domain to active Ras1 molecules by generating constructs linking GAP^Gap1^ to one or three copies of the Byr2 RBD (RBD-GAP^Gap1^ and RasAct-GAP^Gap1^), and expressed these as above in *gap1Δ* cells, at similar levels as GAP^Gap1^ (Figure S4A-B). Remarkably, enhancing GAP^Gap1^ recruitment to Ras1-GTP suppressed the mating defects of *gap1Δ* cells (Figure 7B). This suppression relied on the GAP activity of the fusion constructs (Figure S4C-D). Suppression was partial for RBD-GAP^Gap1^, but complete for RasAct-GAP^Gap1^, which was noticeably recruited to the fusion site (Figure 7A-B), suggesting that the strength of the negative feedback is important to tune the appropriate cellular response.

Finally, we aimed to boost local Ras1-GTP hydrolysis by using the low nanomolar affinity between GFP and GBP (GFP Binding Protein) (Rothbauer et al., 2008) to recruit GAP^Gap1^-GFP directly to the fusion focus labeled with Myo52-GBP-mCherry. In this case, all detectable GAP^Gap1^ localized to sites of polarity (Figure 7C). This locally-recruited GAP efficiently suppressed the lysis of *gap1Δ* cells, though it did not fully restore fusion ability (Figure 7D), as these cells showed an extended fusion reaction, as measured by the duration of the Myo52-labelled fusion focus (Figure 7E). Interestingly, recruitment of GAP^Gap1^ to the fusion focus in otherwise *gap1+* cells caused dominant effects, also significantly extending the duration of the fusion reaction (Figure 7E). This suggests that an exacerbated local Ras1-GTP inhibition delays fusion. Together, these results indicate that the negative feedback that Gap1 provides on Ras1 activity is critical to coordinate the process of cell fusion with cell-cell contact.

## Discussion

The Ras-MAPK cascade is widely used to underlie cell decisions, between proliferation and differentiation, or between life and death (Rauch et al., 2016). Here, we show that a negative feedback regulation of Ras1 activity, promoting cycling between Ras1-GTP and Ras1-GDP, forms an inherent part of the pheromone sensing system in yeast, which serves to mediate both pairing of sexual partners and cell-cell fusion. We propose that the coupling of this feedback inhibition with previously established positive feedback forms an integrated pheromone sensing system that coordinates cell polarization and fusion decisions.

The positive feedbacks that underlie the fission yeast sexual reproduction cycle are well established. At the transcriptional level, pheromone signaling promotes its own expression, such that two cells of opposite mating types stimulate each other (reviewed in (Merlini et al., 2013)). At the cortex, pheromone signaling is coupled to cell polarization, with pheromone release occurring at polarity sites and pheromone sensing promoting polarity site stabilization (Merlini et al., 2016). This forms a positive feedback system through which an initially unstable Cdc42 patch (Bendezu and Martin, 2013) becomes stabilized for cell-cell pairing. This positive feedback likely also occurs in other organisms. For instance in budding yeast, localization of pheromone receptors and transporters is polarized at the projection tip (Ayscough and Drubin, 1998; Kuchler et al., 1993); reciprocally, pheromone receptor activation promotes cell polarization in the direction of the pheromone gradient (reviewed in (Merlini et al., 2013)). Positive feedback between signaling and polarization is further enhanced in preparation for cell fusion, as the actin cytoskeleton forms a concentrated fusion focus, which is immobilized in response to local pheromone-MAPK signaling (Dudin et al., 2015; Dudin et al., 2016). Because the fusion focus is a site of cell wall digestive enzyme release (Dudin et al., 2015), its immobilization leads to local cell wall digestion, a process essential for fusion with a partner cell, but lethal if premature. One characteristic of positive feedbacks is their ability to act as switches, rapidly entraining the system from the off to the all-on situation (Ferrell, 2002). In the case of cell mating, this would have catastrophic lethal consequences because premature cell wall digestion before cell-cell engagement would lead to cell lysis, as is for instance observed upon artificial engagement of the positive feedback in autocrine cells (Dudin et al., 2016). We discuss below how our data support a model in which the positive feedback is adaptively counteracted by Gap1 negative feedback, thus allowing pheromone gradient sensing to consecutively drive dynamic polarization for cell-cell pairing and signal stabilization for cell-cell fusion.

### Gap1 restrains the mating process

Several pieces of evidence indicate that Gap1 negatively regulates mating. First, previous data showed that *gap1Δ* cells exhibit higher transcriptional output upon pheromone stimulation (Weston et al., 2013). Second, we previously showed that Gap1 is necessary to destabilize polarity zones during the cell-cell pairing process and to raise the pheromone sensing threshold above which cells stabilize a zone of polarized growth (Merlini et al., 2016). Consistently, Ras1 in its active form co-localizes with active Cdc42 at exploratory polarity sites, and Gap1 is similarly recruited to these sites. Third, we show here that *gap1Δ* cells mount a premature fusion response leading to frequent lysis. This lysis is not due to ‘unsustainable elongation from multiple tips’ as previously proposed (Weston et al., 2013), as *gap1Δ* cells grow only from one cell pole at any given time (Movie S4). Instead the lysis (but not the excessive growth) is fully suppressed by deleting *fus1*, and thus results from the premature assembly of the fusion focus, leading to local cell wall digestion in absence of a partner cell. This phenotype resembles that of autocrine cells (Dudin et al., 2016), suggesting that *gap1Δ* cells display unrestrained positive feedback. As autocrine *gap1Δ* cells lyse more frequently than autocrine *gap1+* cells, Gap1 likely tempers the positive feedback even in these cells. We conclude that Gap1 acts to dampen pheromone-dependent responses.

### Gap1 forms a negative feedback on Ras1 activity

Transduction of the pheromone signal requires Ras signaling (Nielsen et al., 1992). The molecular connections between the activated pheromone receptor and Ras1 activation are not fully established, though this involves the GEF Ste6 (Hughes et al., 1990; Hughes et al., 1994). In turn, Ras1 activates both the pheromone MAPK cascade by binding directly to the MAP3K Byr2 (Masuda et al., 1995) and Cdc42 by binding the Cdc42 GEF Scd1 (Chang et al., 1994). Consistent with previous genetic analysis (Imai et al., 1991), our data shows that Gap1 functions as a GTPase activating protein for Ras1 both *in vitro* and *in vivo*. Indeed, its deletion leads to excessive Ras1-GTP levels and phenotypes indistinguishable from those of GTP-locked Ras1 alleles. Thus, Gap1 promotes Ras1 cycling between its GTP- and GDP-bound states, which, in addition to its activity *per se*, is important to coordinate the progression of the mating response.

Beyond promoting Ras1-GTP and -GDP cycling, our data show that Gap1 forms a negative feedback. Indeed, Gap1 is recruited to sites of Ras1-GTP both during the vegetative and sexual lifecycle. Though this localization may have been expected from the known interaction of GAP domains with the GTP-bound form of their cognate small GTPase, the Gap1 GAP domain was only poorly recruited to Ras1-GTP sites. Our data indicate that at least two elements in Gap1, its GAP domain and other regions, contribute to its recruitment to Ras1-GTP sites, either through direct Ras1-GTP binding or indirectly by binding downstream components. In addition, the extended duration of Gap1 cortical localization beyond disassembly of dynamic polarity patches and its broader distribution around stable fusion foci suggest that Gap1 localization is not simply reflecting direct Ras1-GTP binding. Upon Ras1-GTP hydrolysis, Gap1 may be handed over to other binding partners at the membrane, or simply remain bound to Ras1-GDP. While further dissection will be required to distinguish between these possibilities, we conclude that Ras1 activity recruits its own inhibitor, thus forming a negative feedback critical for early polarization, cell-cell pairing and coordination of fusion with cell-cell contact.

### Role of negative feedback in cell polarization for cell-cell pairing

Dynamic zones of polarity, as observed early during mating (Bendezu and Martin, 2013), require negative feedback regulation (Martin, 2015; Wu and Lew, 2013). While the molecular mechanisms underlying the formation of zones of Cdc42 activity are not fully elucidated, these likely rely on both spontaneous symmetry-breaking mechanisms, well described in other cells and conditions (reviewed in (Johnson et al., 2011; Martin, 2015; Slaughter et al., 2009)), and coupling to activated pheromone receptors. Ras1-dependent activation of the Cdc42 GEF Scd1 likely serves to couple local pheromone sensing with cell polarization. One interpretation of the outgrowth of *gap1Δ* cell projection at inappropriate location (this work; (Merlini et al., 2016; Weston et al., 2013)) is that these cells do not sense pheromone gradients and break symmetry at random location. The Gap1-dependent transient nature of Ras1 activation at sites of pheromone receptor engagement may impose a local bias to the spontaneous symmetry-breaking mechanisms governing Cdc42 activation. Such mechanism is similar to that envisaged for bud site selection in budding yeast, where the cycling of the Ras-like protein Rsr1 between GTP and GDP forms, couples Cdc42 to spatial landmarks (Bi and Park, 2012). Thus, the cycling between Ras1-GTP and Ras1-GDP, by preventing the progressive activation of the entire cellular Ras1 pool, may allow dynamic coupling of Cdc42 activation to pheromone-engaged receptors.

The ability of cells to pair with a partner appears to scale with the strength at which the Ras1 GAP is recruited to sites of activation. Indeed, we observed a progressively higher frequency of cell pairs upon absence of Gap1, expression of the sole GAP^Gap1^ domain, addition of one RBD or three RBDs to GAP^Gap1^. This indicates that a sufficiently strong negative feedback is necessary to destabilize all the polarity sites that are not stabilized by a reciprocally polarized partner cell.

The Ras1 negative feedback shown here is in line with several negative feedbacks proposed and/or demonstrated to promote spatio-temporal oscillations in other polarized systems, across eukaryotes. These include oscillations in Cdc42 GTPase activity during yeast polarization (Das et al., 2012; Howell et al., 2012; Lee et al., 2015), for which negative feedback was shown to buffer the system and enhance robustness (Howell et al., 2012; Kuo et al., 2014). Additional examples include the growth-mediated recruitment of a Rho GAP in pollen tubes to promote oscillatory growth dynamics (Hwang et al., 2005); or negative feedbacks that keep migratory cells such as *Dictyostelium* in an excitable state (Devreotes and Horwitz, 2015). The parallel with *Dictyostelium* migration is particularly interesting because activation of RasG, one of the 9 Ras homologues in this organism, is one of the earliest polarized responses to the cAMP chemoattractant (Sasaki et al., 2004). In addition, deletion of its GAP DdNF1 leads to near homogeneous RasG activation at the cell cortex and loss of directional sensing (Zhang et al., 2008), similar to the case of *gap1Δ* in *S. pombe*. Whether this GAP is part of negative feedback or contributes to global RasG-GTP hydrolysis is unknown, but it is intriguing that its mammalian homologue, the Neurofibromin NF1, has been proposed to function in Ras feedback inhibition (Hennig et al., 2016). These parallels suggest that feedback regulation of Ras activity contributes to spatial patterning across eukaryotic species.

### Local high MAPK signaling at the fusion site

One particularity of Ras is its position at crossroads between polarity and MAPK signaling regulation. Ras1-dependent activation of the MAPK cascade and its recruitment to the fusion focus is likely to be a critical event for cell fusion. We previously showed that forced recruitment of the MAP2K Byr1 to the fusion focus is sufficient to stabilize the structure and induce fusion attempts (Dudin et al., 2016). Our data now show that constitutive Ras1 activation promotes Byr1 localization to the fusion focus even in absence of pheromone receptor localization. Although this is consistent with previous observation that active Ras1 recruits the MAP3K Byr2 to the cell cortex (Bauman et al., 1998), we note that, because both pheromone receptor and Ras1 are active on a broad cortical zone in this case, additional inputs must restrict the specific localization of Byr1 on the focus. Additive phenotypes of mutants in the pheromone receptor-coupled Gα *gpa1* and *ras1* (Xu et al., 1994) suggest Gpa1 may play a role. Previous observations that *scd1* deletion suppresses cell lysis of *gap1Δ* cells (Weston et al., 2013) and that Byr1 focal localization requires Fus1 (Dudin et al., 2016) similarly suggest a role for Cdc42 and the actin fusion focus.

Interestingly, recent work showed that elevated MAPK signaling also likely promotes cell fusion in *S. cerevisiae* (Conlon et al., 2016). This is noteworthy particularly because the molecular wiring of the pheromone-MAPK pathways differs significantly in the two yeasts, with the *S. cerevisiae* pheromone-MAPK cascade signaling independently of Ras and using a MAPK scaffold, Ste5, absent in *S. pombe*. In *S. cerevisiae*, the MAPK cascade also accumulates at the shmoo tip through mechanisms that involve direct transport of Ste5 scaffold along formin-nucleated actin cables (Qi and Elion, 2005), enzyme-substrate interactions (Maeder et al., 2007) or binding to the polarisome component Spa2 (Sheu et al., 1998). This suggests that, despite the distinct molecular pattern, the concentration of the MAPK cascade at the fusion site may be a common switch decision to induce cell fusion.

### Role of negative feedback in coordinating fusion with cell-cell contact

We have shown that negative feedback on Ras1 activity is essential to prevent premature fusion events. Expression of GAP^Gap1^ domain alone was unable to prevent premature fusion attempts. By contrast, engineering recruitment of the GAP^Gap1^ to Ras1-GTP with either one or three RBDs restored, partly or fully, respectively, the coordination between cell contact and fusion. The finding that GAP^Gap1^ domain promoted pair formation but did not protect against lysis further suggests that a stronger negative feedback is required to couple fusion with cell contact than to promote cell pair formation. Interestingly, engineering an even higher local concentration of GAP^Gap1^ at the fusion site, by forced recruitment to Myo52-GBP in addition to endogenously expressed Gap1, had only minor effects: while it significantly delayed fusion, it had no effect on the overall fusion efficiency. We conclude that a strong negative feedback that promotes rapid Ras1-GTP hydrolysis is key to coordinate consecutive responses to pheromone gradient signaling.

Negative feedbacks have been shown to act at multiple levels of the budding yeast pheromone signaling pathway, which has been extensively used to dissect MAPK cascades (Atay and Skotheim, 2017). Interestingly, eliminating any single one of these negative feedbacks did not significantly alter the graded transcriptional response (Takahashi and Pryciak, 2008). However, a negative feedback involving MAPK Fus3 activity and the regulator of G-protein signaling Sst2 (a GAP for the receptor-coupled Gα) was shown to help align the dose-response between pheromone receptor occupancy and cellular outputs (Yu et al., 2008). Though effects of negative feedback on cell fusion, which is an all-or-none switch-like output of the MAPK signal, have to our knowledge not been investigated in *S. cerevisiae*, our results are in line with this idea that Gap1 serves to ensure the physiological polarization and fusion responses are aligned with appropriately high pheromone exposures. Because the *S. pombe* pheromone-Ras-MAPK cascade is very closely related to the growth factor-Ras-ERK mammalian cascade (Hughes et al., 1993), similar feedbacks may operate to align physiological responses in mammalian cells.

### Mating progression through integrated positive and negative feedbacks

As Gap1-dependent feedback blocks premature polarization and fusion attempts, one question is what provides order to the mating process. We propose that the physical distance between two partner cells inherently regulates the strength of the pheromone-dependent positive feedback signal and thus orders morphological responses. In support for this idea, pheromones are released locally at sites of polarity (Merlini et al., 2016), and signal perception promotes local polarization, which leads to local secretion for growth. Thus, as the proximity between cells increases, the site of pheromone release narrows, with the consequence that the profile of the perceived pheromone gradient becomes progressively sharper and its amplitude higher. Consistently, Ras1-GTP shows significantly narrower distribution and higher peak intensity at fusion time than during cell pairing. As the polarity site narrows to a tight focus, the local concentration of secreted cell wall hydrolases dominates that of cell wall synthases leading to local cell wall digestion (Dudin et al., 2015). The negative feedback, by destabilizing the polarity site, extends the range of pheromone concentrations supporting dynamic polarization, while still allowing high pheromone concentrations to trigger cell-cell fusion. Thus one way to think about the Gap1-dependent negative feedback is as setting a cell-to-cell distance threshold to ensure that fusion focus stabilization happens only once the local pheromone signal is high enough, and thus couple fusion with cell-cell contact.

## Experimental procedures

### Strains, Media, and Growth Conditions

Strains used in this study are listed in Table S1. Standard genetic manipulation methods for *S. pombe* and *S. cerevisiae* transformation and tetrad dissection were used. For microscopy of fission yeast cells during exponential growth, cells were grown in Edinburgh minimal medium (EMM) supplemented with amino acids as required. For biochemistry experiments, cells were grown in rich Yeast extract medium (YE) or minimal sporulation medium with nitrogen (MSL+N). For assessing cells during the mating process, liquid or agar minimal sporulation medium without nitrogen (MSL-N) was used (Egel et al., 1994; Vjestica et al., 2016). All live-cell imaging during sexual lifecycle was performed on MSL-N agarose pads (Vjestica et al., 2016). Mating assays were performed as in (Vjestica et al., 2016). For microscopy of budding yeast cells, cells were grown in Synthetic Defined medium (SD) supplemented with amino acids as required.

Gene tagging was performed at endogenous genomic locus at the 3’ end, yielding C-terminally tagged proteins, as described (Bähler et al., 1998). Tagging with sfGFP was performed as in (Dudin et al., 2015). Tagging with GBP-mCherry was performed as in (Dudin et al., 2016). N-terminal tagging of Ras1 with GFP was performed as in (Merlini et al., 2016). Gene deletion was performed as described (Bähler et al., 1998). Gene tagging and deletion were confirmed by diagnostic PCR for both sides of the gene.

Construction of fission yeast strains expressing RasAct^GFP^ (3x-Byr2^RBD^-3xGFP) was done by integration of RasAct^GFP^ under *pak1* promoter at the *leu1+* locus. A fragment encoding the RBD domain of Byr2 (Fig. 2A) was cloned in 3 tandem repeats into pAV49 (pJK210, a kind gift from Dr. Aleksandar Vjestica, UNIL): first, Byr2-RBD was amplified from genomic DNA with primers osm2930 (5′-tccc<u>cccggg</u>CGAGAGTTTCCACGTCCATG) and osm2932 (5′-cg<u>ggatcc</u>AGGAGAAAGGGAGGACTGTG), digested with XmaI and BamHI and ligated to similarly treated pAV49 to generate plasmid pJK210-*1x-byr2-RBD*, second, Byr2-RBD was amplified from genomic DNA with primers osm2933 (5′- cg<u>ggatcc</u>CGAGAGTTTCCACGTCCATG) and osm2934 (5′- gc<u>tctaga</u>AGGAGAAAGGGAGGACTGTG), digested with BamHI and XbaI and ligated to similarly treated pJK210-*1x-byr2-RBD* to generate plasmid pJK210-*2x-byr2-RBD*, third, Byr2-RBD was amplified from genomic DNA with primers osm2935 (5′-gc<u>tctaga</u>CGAGAGTTTCCACGTCCATG) and osm3082 (5′-tcc<u>ccgcggTTAATTAA</u>AGGAGAAAGGGAGGACTGTG), digested with XbaI and SacII and ligated to similarly treated pJK210-*2x-byr2-RBD* to generate plasmid pJK210-*3x-byr2-RBD* (pSM1627). The *3x-byr2-RBD* fragment was then excised from plasmid pSM1627 through digestion with XmaI and PacI and ligated into similarly treated pAV55 (a pJK148-based vector containing the *pak1* promoter, a kind gift from Dr. Aleksandar Vjestica, UNIL) to generate plasmid pINT-*Ppak1-3x-byr2-RBD-3xGFP-kanMX-leu1+* (pSM1628). Finally, pSM1628 digested with AfeI was stably integrated as a single copy at the *leu1+* locus in the yeast genome. In primer sequences, restriction sites are underlined.

Construction of budding yeast strains expressing RasAct^GFP^ was done by integration of RasAct^GFP^ under *pRPL24A* promoter at the *ura3* locus. First, RasAct^GFP^ was amplified from pSM1628 with primers osm3900 (5′- gg<u>actagt</u>ATGGGAGGTCCCGGGCGA) and osm3901 (5′- acgc<u>gtcgac</u>GGCGCGCCGATATTAAAG), digested with SpeI and SalI and ligated to similarly treated pSM1890 (pRS413, a gift from Dr. Simonetta Piatti) to generate plasmid pSM1898 (pRS413-*Pgpd-RasAct^GFP^-Tcyc1*), second, the *RasAct^GFP^-Tcyc1* fragment was excised from plasmid pSM1898 through digestion with SpeI and KpnI and ligated into similarly treated pSM1974 (pDA133-*pRPL24A*, a gift from Dr. Serge Pelet) to generate plasmid pSM1977 (*pRPL24A- RasAct^GFP^-Tcyc1*). Finally, pSM1977 digested with BstBI was stably integrated as a single copy at the *ura3* locus in the yeast genome. In primer sequences, restriction sites are underlined.

Construction of strains expressing the constitutively active *ras1^G17V^* and *ras1^Q66L^* or the GDP-bound *ras1^S22N^* alleles was done by integration at the endogenous *ras1* locus. First, a fragment including ras1 coding region, 5′- and 3′- extensions was amplified from genomic DNA with primers osm2148 (5′-acgc<u>gtcgac</u>CACATTTTAACGAGCTTAAGACC) and osm2149 (5′-tcc<u>cccggg</u>GCTGCTAATAATTGTGTTAAATG), digested with SalI and XmaI and ligated to similarly treated pSM1232 to generate plasmid pSM1316 (pSP72-*ras1*). Second pSM1316 was subjected to site directed mutagenesis with primers osm2163 (5′-GGTAGTTGTAGGAGAT**GTT**GGTGTTGGTAAAAGTG) and osm2164 (5′-

CACTTTTACCAACACC**AAC**ATCTCCTACAACTACC) to generate plasmid pSM1320 (pSP72-*ras1^G17V^*), with primers osm2167 (5′-

GTATTGGACACGGCCGGT**CTA**GAGGAATATTCCGCTATG) and osm2168

(5′- CATAGCGGAATATTCCTC**TAG**ACCGGCCGTGTCCAATAC) to generate plasmid pSM1322 (pSP72-*ras1^Q66L^*) or with primers osm2165 (5′-

GGTGGTGTTGGTAAA**AAT**GCTTTGACAATTCAAT) and osm2166 (5′-

ATTGAATTGTCAAAGC**ATT**TTTACCAACACCACC) to generate plasmid pSM1321 (pSP72- *ras1^S22N^*). Finally, pSM1320, pSM1321 or pSM1322 digested with SalI and XmaI were stably integrated as single copy at the *ras1* locus in the yeast genome, through transformation of a *ras1::ura4+* strain and selection on agar plates containing 5-Fluoroorotic Acid (5-FOA). In primer sequences, restriction sites are underlined, inserted mutations are bold.

Construction of strains expressing the *gap1^R340A^* or *gap1^R195AR340A^* alleles was done by integration at the endogenous *gap1* locus. First, a fragment including gap1 coding region, 5′- and 3′- extension was amplified from genomic DNA with primers osm2232 (5′-acgc<u>gtcgac</u>CTTAGTATAATATCCATCCTTG) and osm2233 (5′-tcc<u>cccggg</u>GACGATTAATGTATAAGAAAC), digested with SalI and XmaI and ligated to similarly treated pSM1232 to generate plasmid pSM1348 (pSP72-*gap1*). Second pSM1348 was subjected to site directed mutagenesis with primers osm2567 (5′-

GATTTTTCTTTCTT**GCT**TTCGTTAATCCAGC) and osm2568 (5′-

GCTGGATTAACGAA**AGC**AAGAAAGAAAAATC) to generate plasmid pSM1513 (pSP72-*gap1^R340A^*). Third, pSM1513 was subjected to site directed mutagenesis with primers osm2571 (5′- GTTTTGTCTCTGCTT**GCG**GCTAATACTCCGG) and osm2572 (5′- CCGGAGTATTAGC**CGC**AAGCAGAGACAAAAC) to generate plasmid pSM1645 (pSP72-*gap1 ^R195AR340A^*). Finally, pSM1513 or pSM1645 digested with SalI and XmaI were stably integrated as single copy at the *gap1* locus in the yeast genome, through transformation of a *gap1::ura4+* strain and 5-FOA selection. In primer sequences, restriction sites are underlined, inserted mutations are bold.

Construction of strains in Figure 5 expressing Gap1^FL^, GAP^Gap1^ or the fusion constructs RBD-GAP^Gap1^ and RasAct-GAP^Gap1^ was done by integration under *pak1* promoter at the *leu1+* locus. Gap1 gene was amplified from genomic DNA with primers osm4122 (5′- tccc<u>cccggg</u>ACTAAGCGGCACTCTGGTACCC) and osm4123 (5′-cc<u>ttaattaa</u>CTTTCGTAAAAACAATTGTTC), digested with XmaI and PacI and ligated to similarly treated pSM1628 to generate plasmid pSM1984 (pINT- *Ppak1-*Gap1^FL^-3XGFP*-kanMX-leu1+*). The GAP domain of Gap1 was amplified from genomic DNA with primers osm3922 (5′- tccc<u>cccggg</u>CTTCAGTTGTATGGAGCGTTG) and osm3921 (5′-cc<u>ttaattaa</u>TAAATCAGGAATTGAAGAATCCCATCG), digested with XmaI and PacI and ligated to similarly treated pSM1628 to generate plasmid pSM1954 (pINT-*Ppak1-* GAP^Gap1^-3XGFP*-kanMX-leu1+*). The GAP domain of Gap1 was amplified from genomic DNA with primers osm3922 and osm3923 (5′-tccc<u>cccggg</u>TAAATCAGGAATTGAAGAATCCCATCG) digested with XmaI and ligated to similarly treated pSM1628 to generate plasmid pSM1896 (pINT-*Ppak1-* RasAct-GAP^Gap1^*-kanMX-leu1+*) or pSM1603 (pINT-*Ppak1-1xbyr2-RBD-3xGFP-kanMX-leu1+*) to generate plasmid pSM1897 (pINT-*Ppak1-*RBD-GAP^Gap1^*-kanMX-leu1+*). Plasmids carrying mutations in the Gap domain of Gap1 were generated by site directed mutagenesis as explained in the previous paragraph. Finally, plasmids digested with AfeI were stably integrated as a single copy at the *leu1+* locus in the yeast genome. In primer sequences, restriction sites are underlined.

For recombinant protein production, *ras1* was amplified from a cDNA library with primers osm1941 (5′-cg<u>ggatcc</u>ATGAGGTCTACCTACTTAAGAGAGTAC) and osm1942 (5′-ccg<u>ctcgag</u>CTAACATATAACACAACATTTAGTTG) digested with BamHI and XhoI and ligated to similarly treated pSM394 (pGEX-4T-1) to generate plasmid pSM1282 (pGEX-4T-1-*ras1*). RBD fragment was amplified from genomic DNA with primers osm2933 and osm3232 (5′-tccc<u>cccggg</u>**TTA**AGGAGAAAGGGAGGACTGTG) digested with BamHI and XmaI and ligated to similarly treated pSM394 to generate plasmid pSM1713 (pGEX-4T-1-*RBD*). *gap1* was amplified from a cDNA library with primers osm2152 (5′-aaggaaaaaa<u>gcggccgc</u>cATGCTTCAGTTGTATGGAGCGTTG) and osm2311 (5′-ccg<u>ctcgag</u>TTACTTTCGTAAAAACAATTGTTC) digested with NotI and XhoI and ligated to similarly treated pSM819 (pMAL-TEV) to generate plasmid pSM1401 (pMAL-TEV-*gap1*). Note that we were not able to purify the full length Gap1 recombinant protein and a protein lacking the first 116 aa was used for the *in vitro* GAP experiments. In primer sequences, restriction sites are underlined, stop codon is bold.

### Mating Assays

Mating assays were performed as in (Vjestica et al., 2016). Briefly, pre-cultures of cells were grown over night in MSL+N at 25°C to reach an OD600 of between 0.5 and 1. Cultures were then diluted to an OD600 of 0.025 in MSL+N and grown for 16-20 hours to an OD600 of between 0.5 and 1 at 30°C. Cells were washed three times with MSL-N, diluted to an OD600 of 1.5 in 1 ml MSL-N and incubated at 30°C for 1-4 h (depending on the mating stage to be visualized). Cells were mounted onto MSL-N agarose pads (2% agarose) before imaging in overnight movies or incubated at 18°C or at 25°C overnight before imaging. Fusion efficiency, mating efficiency and percentage of cell lysis were measured as in (Dudin et al., 2015; Dudin et al., 2016). For pheromone treatments, P-factor pheromone was purchased from Pepnome (Zhuhai City, China) and used from a stock solution of 1 mg/ml in methanol. M-factor was synthetized and purchased from Schafer-N and used from a stock solution of 2mg/ml in methanol. Different concentrations of pheromones were directly added to the melted MSL-N agarose before mounting cells on the pads. Cells were then imaged overnight or incubated at 25°C prior to imaging. Methanol was used as a control.

### Microscopy and Image Analysis

The DeltaVision platform (Applied Precision) described previously (Bendezu and Martin, 2011) was used for time-lapse imaging overnight, quantitative analyses of mating efficiency, fusion efficiency and cell lysis, short time-lapse imaging and quantification of cortical signal (Figures 1, 3G-I, 4C-F, 5B, 6C, 7B-E, S2, S3, S4C). Spinning disk microscopy previously described (Bendezu and Martin, 2011; Dudin et al., 2015) was used for high-temporal resolution and Z-stack maximal projection images, time-lapse imaging and quantification of cortical signal (Figures 2B-F, 3A-F, 4F, 6A, B, E, 7A, S1, S4D). Kymographs in Figures 1, 3, 6 and S2 were constructed in ImageJ version 1.47 (NIH) by drawing a 3-pixel-wide line at the cell tip or around the cell cortex.

Quantification of cortical fluorescence at the cell tip in Figure 2D and whole cortical fluorescence in Figure 2F was done by using the sum projection of five consecutive images. The intensity of a 3 pixel wide segment was collected. Images were corrected for the external background. Quantification of cortical fluorescence at the shmoo or fusion tip in Figures 3G and 3H was done by drawing a 3-pixel-wide line at the cell tip. The curves in Figure 3G were centered on the maximum pixel values for the Myo52-tdTomato channel (not shown in the figure). Quantifications in Figure 3H were done on the 5 brightest pixels in the selected area. Images were corrected for the external background. Quantification of RasAct^GFP^ fluorescence over time during the fusion process in Figure 3I was done by drawing a 3-pixel-wide line at the cell tip. The curves were aligned to the time of fusion visualized by observing the diffusion of a cytosolic marker from one cell to the other (Vjestica et al., 2016). Images were corrected for the internal background by substracting the average intensity of a reference cell in the same field at each timepoint. A Lilliefors test was used to check that data were not significantly different from normal. A t-test with Bonferroni correction for multiple comparisons was used for statistical analysis. Quantification of Gap1-GFP, Ste6-sfGFP and RasAct^GFP^ fluorescence at the fusion focus in Figure 6F was done by drawing a 3-pixel-wide line at the fusion site. The curves were centered on the maximum pixel values for the Myo52-tdTomato channel (expressed in each strain). Images were corrected for the external background.

To analyze the lifetime of the Scd2-mCherry/Gap1-GFP patch during exploration in Figure 6C-D, *h- sxa2Δ* cells co-expressing Gap1-GFP and Scd2-mCherry were imaged in the presence of 0.01 μg/ml P-factor (to promote exploration, but not stabilization of the patch; (Bendezu and Martin, 2013)) every 30 seconds for 10 minutes. To follow the entire processes of Scd2-mCherry/Gap1-GFP patch formation and disassembly we selected cells in which the Scd2-mCherry patch formed within the first half of the movie. We further selected cells in which the exploring patch did not re-appear at the same location. To measure photo-bleaching, we fitted an exponential decay function with decay constant *r*_PB_ to the whole cell signal after subtracting the out-of-cell background. To correct for photo-bleaching, the pixel intensity at every time point was corrected by multiplying by 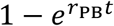 after subtracting the out-of-cell background. For some of the movies in which the slide was drifting, the translation function from StackReg plugin for imageJ (Thevenaz et al., 1998) was used to align the slices. The GFP channel was used to manually draw the cell boundary around the cell cortex, before recording the intensity of Gap1-GFP and Scd2-mCherry for all time points. We defined the width of the Scd2 patch as the region over which the Scd2-mCherry signal exceeded the average cortical signal in neighboring regions and obtained average intensities of Gap1-GFP and Scd2-mCherry for this region. Patch width ranged between 11-16 pixels equivalent to 1.4-2.0 μm. We also did the same analysis using the full width at half maximum signal of the Scd2-mCherry, which led to the same conclusions. Profiles were normalized, after subtraction of the value in a neighboring zone of similar size, to the maximal value at the patch. Examples of such traces are shown in Figure S3B-D. In order to plot an average of these traces, we aligned the Scd2-mCherry intensity traces using the continuous-alignment method of (Berro and Pollard, 2014), and co-aligned the Gap1-GFP traces.

### Biochemistry methods

Recombinant proteins were produced in BL21 cells and purified on GST-Sepharose (GE Healthcare) or amylose beads (New England Biolabs). Cells were grown overnight in 10ml LB-Amp (LB with 100μg/ml ampicillin) at 37°C. The day after 6.25ml of the saturated culture were inoculated in 250ml of LB-Amp, grown 3 hours at 37°C and cooled down 15 min at 4°C. Protein expression was induced by the addition of 100μM IPTG for 5-6 hours at 18°C. For purification of MBP-Gap1 and MBP-Gap1^R340A^, bacterial pellets were resuspended in 10ml cold resuspension buffer (50mM Tris-HCl, pH=8, 1mM EDTA, 100mM KCl, PMSF), sonicated 3 times for 30sec (50% power amplitude), incubated 30min with 1% Triton-X100 at 4°C, and centrifuged 15min at 4°C at 10,000g. Soluble extract was incubated with 400μl of amylose beads for 2h at 4°C. Finally, beads were washed 3 times with cold resuspension buffer and eluted in three steps in 100μl elution buffer (50mM Tris-HCl, pH=8, 1mM EDTA, 100mM KCl, 10mM maltose). For purification of GST-Ras1 and GST-RBD, bacterial pellets were resuspended in 5ml cold PBS 1X containing PMSF), sonicated 6 times for 30sec (40% power amplitude), incubated 30min with 1% Triton-X100 at 4°C, and centrifuged 15min at 4°C at 10,000g. Soluble extract was incubated with 200μl of GST-Sepharose beads for 2h at 4°C. Finally, beads were washed 3 times with 1X PBS and eluted in three steps in 100μl elution buffer (50mM Tris-HCl, pH=8, 15mM reduced glutathione).

For *in vitro* GAP assays (Geymonat et al., 2002) 1 mg of purified GST-Ras1 was resuspended in 70μl of loading buffer (20mM Tris-HCl, pH 7.5, 25mM NaCl, 5mM MgCl_2_, 0.1mM DTT) containing 1.5μl γ -[^32^P]GTP for 10 min at 30°C. After cooling on ice, 10μl were incubated in 50μl reaction buffer (20mM Tris-HCl, pH 7.5, 2mM GTP, 0.6μg/μl BSA) with 18μg of purified MBP-Gap1, MBP-Gap1^R340A^ or MBP. The reactions were incubated at 30°C, and every 5 minutes, 10μl of reaction was diluted in 990μl of cold washing buffer (20mM Tris-HCl, pH 7.5, 50mM NaCl, 5mM MgCl_2_). The samples were filtered through pre-wetted nitrocellulose filters (Millipore), washed three times with 4ml of cold washing buffer and air-dried. The amount of radioactive nucleotide bound to the protein was determined by scintillation counting.

For Ras-GTP pulldown (Soto et al., 2010), extracts from yeast cells grown in 200ml of YE or 200ml MSL-N (pre-growth in MSL+N and shift to MSL-N for 4 hours) were prepared in cold Binding Buffer (25mM TRIS-HCl pH7.5, 1mM DTT, 30mM MgCl_2_, 40mM NaCl, 0.5% NP40) containing Anti Proteolitic (Roche) and phostop (Roche) tablets. Cell lysates were obtained at 4°C via mechanical brakeage with acid treated glass beads (Sigma) in a BeadBeater homogenizer (10 times at 4,5V for 30 sec with 30 sec break on ice every cycle) and centrifugation 20 min at 4°C at 10,000g. 4-6 mg of protein extract was incubated with 20μl of GST-Sepharose beads and 25 μg of purified GST-RBD for 1 hour at 4°C. Beads were washed 3 times with 500μl of cold washing buffer 1 (25mM TRIS-HCl pH7.5, 1mM DTT, 30mM MgCl_2_, 40mM NaCl, 1% NP40) and 2 times with 500μl of cold washing buffer 2 (25mM TRIS-HCl pH7.5, 1mM DTT, 30mM MgCl_2_, 40mM NaCl) containing protease inhibitor cocktail. Proteins were resolved by SDS-PAGE for Western blot analysis.

Standard protocols were used for SDS-PAGE and Western blot analysis. Antibodies used on western blots were: anti-GFP monoclonal antibody (Roche) and anti-TAT1 monoclonal antibody for α-tubulin detection.

Figures were assembled with Adobe Photoshop CS5 and Adobe Illustrator CS5. All error bars are standard deviations. All experiments were done minimum three independent times.

## Acknowledgements

We thank B. Hegemann, S. Pelet, S. Piatti and A. Vjestica for strains and plasmids, S. Mitri for help with statistics, D. Vavylonis for discussions and critical input, and S. Pelet and lab members for critical reading of the manuscript. This work was supported by a Swiss National Science Foundation (31003A_155944) and an ERC Consolidator Grant (CellFusion) to SGM. BK was supported by NIH grant R01GM098430 (to Dimitrios Vavylonis).

## Figure legends

**Figure S1:**
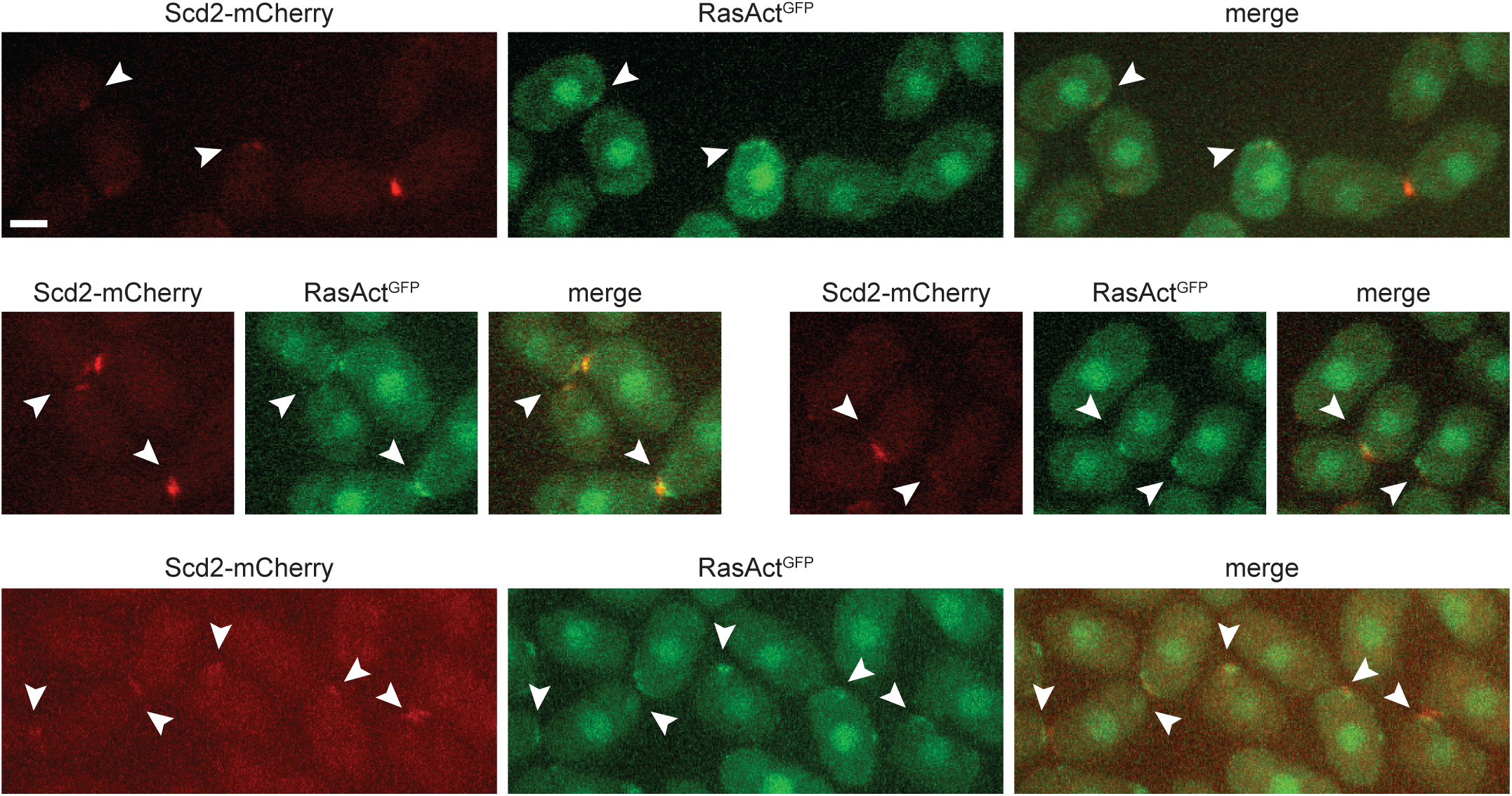
Active Ras1 co-localizes with Scd2 during mating. Spinning disk confocal projections of *h90* wild-type strains showing colocalization of Scd2-mCherry (red) and RasAct^GFP^ (green) during mating. Arrowheads highlight zones of co-localization. Bars = 2 μm.

**Figure S2:**
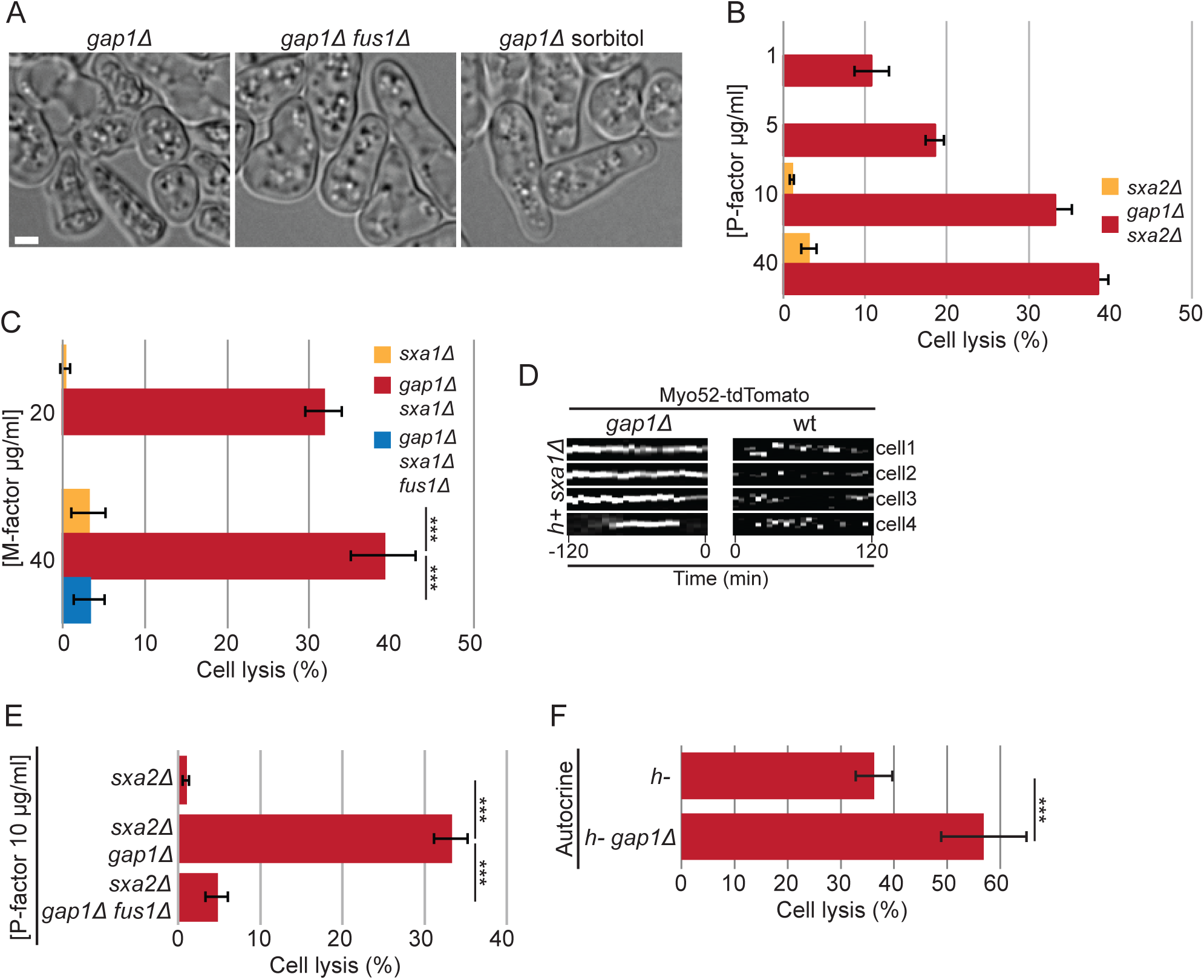
The deletion of Gap1 causes precocious fusion attempts. (A) DIC images of homothallic *gap1*Δ cells treated or not with 1.2M sorbitol or lacking *fus1* (*fus1*Δ) after 14h in MSL-N. (B) Percentage of cell lysis of *h-sxa2*Δ and *h- sxa2Δ gap1*Δ cells, 14h after the addition of synthetic P-factor at indicated concentrations (n > 200 for 3 independent experiments). (C) Percentage of cell lysis of *h+ sxa1*Δ, *h+ sxa1Δ gap1*Δ and *h+ sxa1Δ gap1Δ fus1*Δ cells, 14h after the addition of synthetic M-factor at indicated concentrations (n > 200 for 3 independent experiments). *** indicates p-value^t-test^ ≤ 6.8 x 10^-5^. (D) Kymographs of four representative cell tips showing the formation of a transient Myo52 focus in *h+ sxa1*Δ cells and a stable Myo52 focus in *h+ sxa1Δ gap1*Δ mutants exposed to 20μg/ml M-factor. (E) Percentage of cell lysis of *h- sxa2*Δ, *h- sxa2Δ gap1*Δ and *h- sxa2Δ gap1Δ fus1*Δ cells, 14h after synthetic P-factor addition (n > 500 for 3 independent experiments). *** indicates 5.54 x 10^-6^ ≤ p-value^t-test^ ≤ 1.8 x 10^-5^. (F) Percentage of cell lysis of wt and *gap1*Δ autocrine M-cells (*mam2Δ::map3*; n > 200 for 3 independent experiments). *** indicates p-value^t-test^ ≤ 1.5 x 10^-5^. Bars = 2 μm.

**Figure S3:**
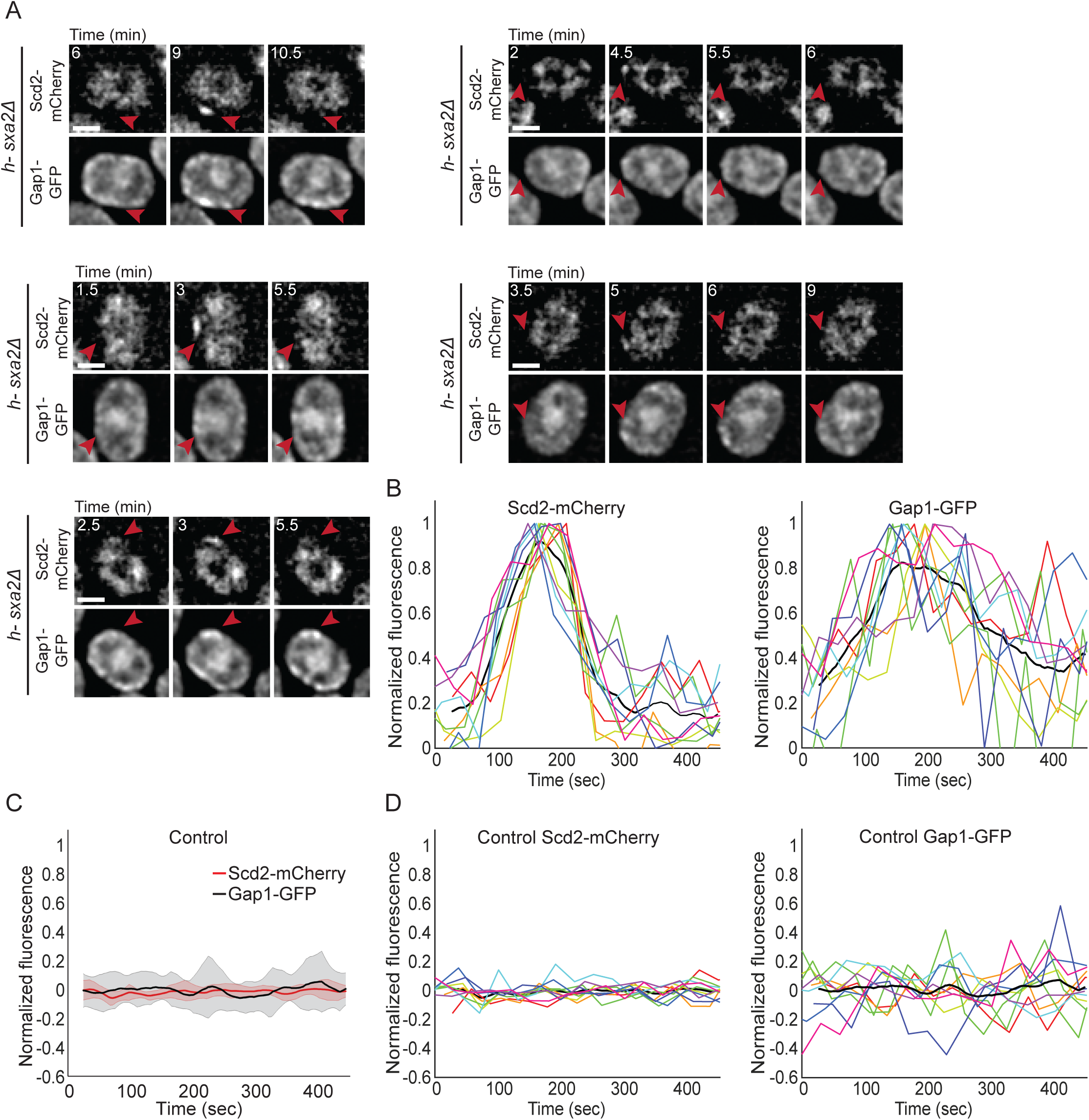
Gap1 disappearance from the exploratory patches is delayed compared to Scd2. (A) Single plane images of *h- sxa2*Δ cells expressing Scd2-mCherry and Gap1-GFP treated with 0.01 μg/ml of synthetic P-factor as in Figure 6C. The red arrowhead highlights the location of one patch of Scd2-mCherry that appears and disappears in the course of the timelapse and the corresponding location in the Gap1-GFP channel. (B) Individual cortical profiles of Scd2-mCherry (left) and Gap1-GFP (right) normalized fluorescence intensity at the exploratory patch in 10 different cells as in (A). The average curves are shown in Figure 6D. (C, D) Control quantification of cortical profiles of Scd2-mCherry and Gap1-GFP normalized fluorescence intensity at a cortical region distinct from the exploratory patch. Panel (C) shows the average and (D) the individual profiles. In (C) the thick line represents the average and the corresponding shaded area the standard deviation. Bars = 2 μm.

**Figure S4:**
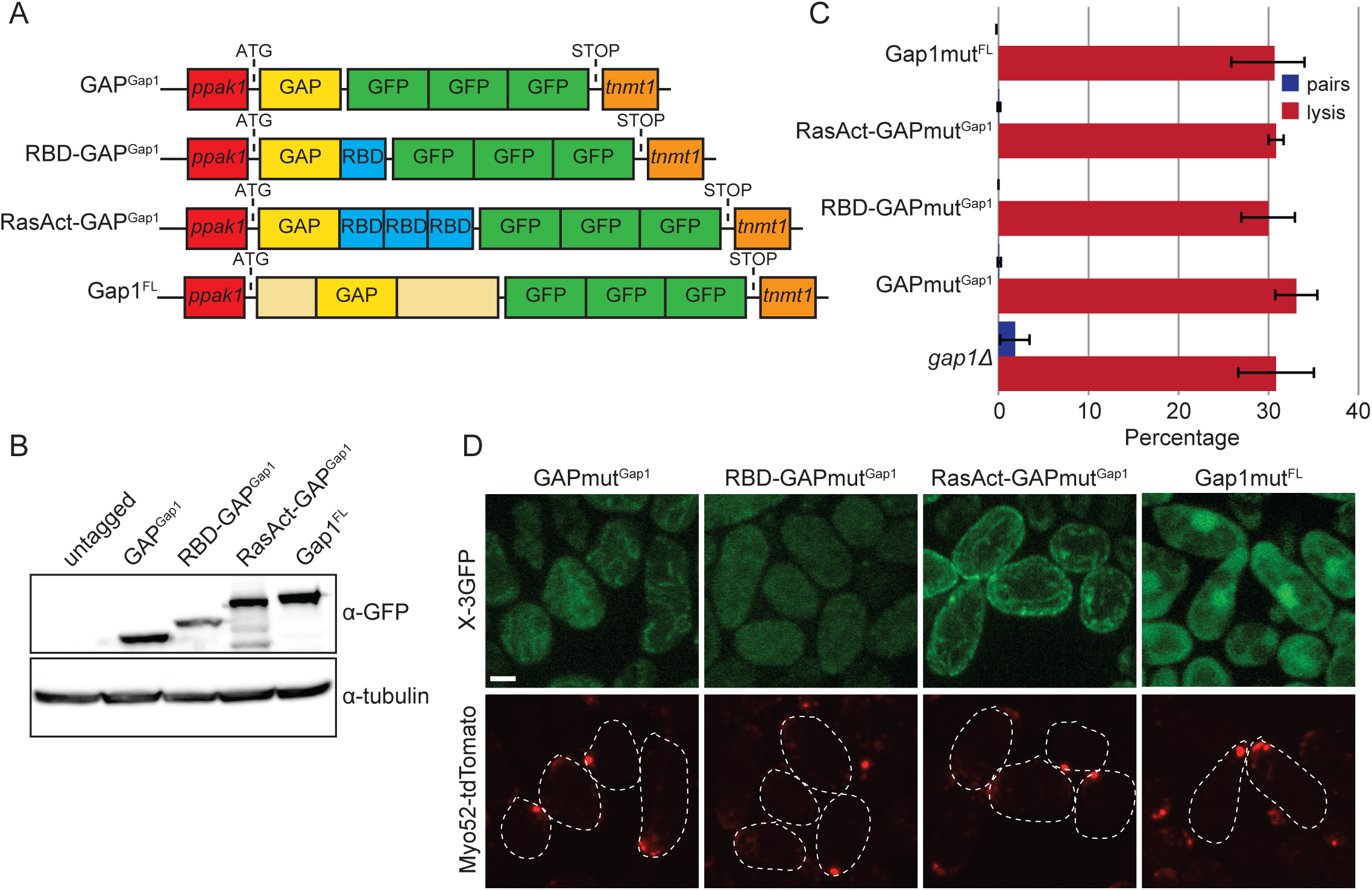
The Gap1 negative feedback requires Gap1 GAP activity to prevent premature fusion attempts. (A) Scheme of the constructs analyzed in Figure 7A-B. All the constructs were expressed from *pak1* promoter, fused to three copy of GFP and integrated at the *ura4+* locus of *gap1*Δ strains expressing Myo52-tdTomato to visualize the fusion focus. (B) Western blot showing similar expression levels of the constructs described in (A). (C) Percentage of cell lysis and cell pair formation of homothallic *gap1*Δ cells expressing or not constructs as described in (A) but carrying point mutations (mut = R195A R340A) in the GAP domain that abolish GAP activity, 12 hours after nitrogen starvation (n > 300 for 3 independent experiments). (D) Representative images of the localization of the mutant constructs as in (C). See Figure 7A for comparison with wildtype constructs. Bars = 2 μm.

**Supplemental Movie1: Fusion focus formation and cell lysis of *ras1^G17V^* cells upon pheromone exposure.** Deconvolved single z-plane epifluorescence and DIC timelapse of *h- sxa2Δ ras1^G17V^* cells expressing Myo52-tdTomato treated with 1 *μ*g/mL P-factor. Cells form mating projections and assemble a stable Myo52 focus. A proportion of cells undergo fusion attempts that result in cell lysis. Note that in some cells the Myo52 signal is out of focus and thus is not visible at the shmoo tip.

**Supplemental Movie2: Premature fusion attempts and lysis of *gap1*Δ cells during mating.** Deconvolved single z-plane epifluorescence and DIC timelapse of *h90 gap1*Δ cells expressing Myo52-tdTomato. Shmooing cells assemble a stable Myo52 focus and attempt precocious fusion thus causing cell lysis. Note that visible extruded cell material at the shmoo tip of the top cell indicates cell lysis at this location.

**Supplemental Movie3: *gap1*Δ cells form an actin fusion focus and engage in untimely fusion during mating.** Deconvolved single z-plane epifluorescence and DIC timelapse of *h90 gap1Δ* cells expressing a GFP-CHD marker for actin visualization. Cells form a stable actin focus at the shmoo tip, with some cells undergoing lysis.

**Supplemental Movie4: *gap1*Δ cells form several consecutive shmoos at inappropriate location.** Deconvolved single z-plane epifluorescence and DIC timelapse of *h90 gap1Δ* cells expressing a GFP-CHD marker for actin visualization. Note that the cell with more than one shmoo grows from a single site at any given time.

**Table S1.**
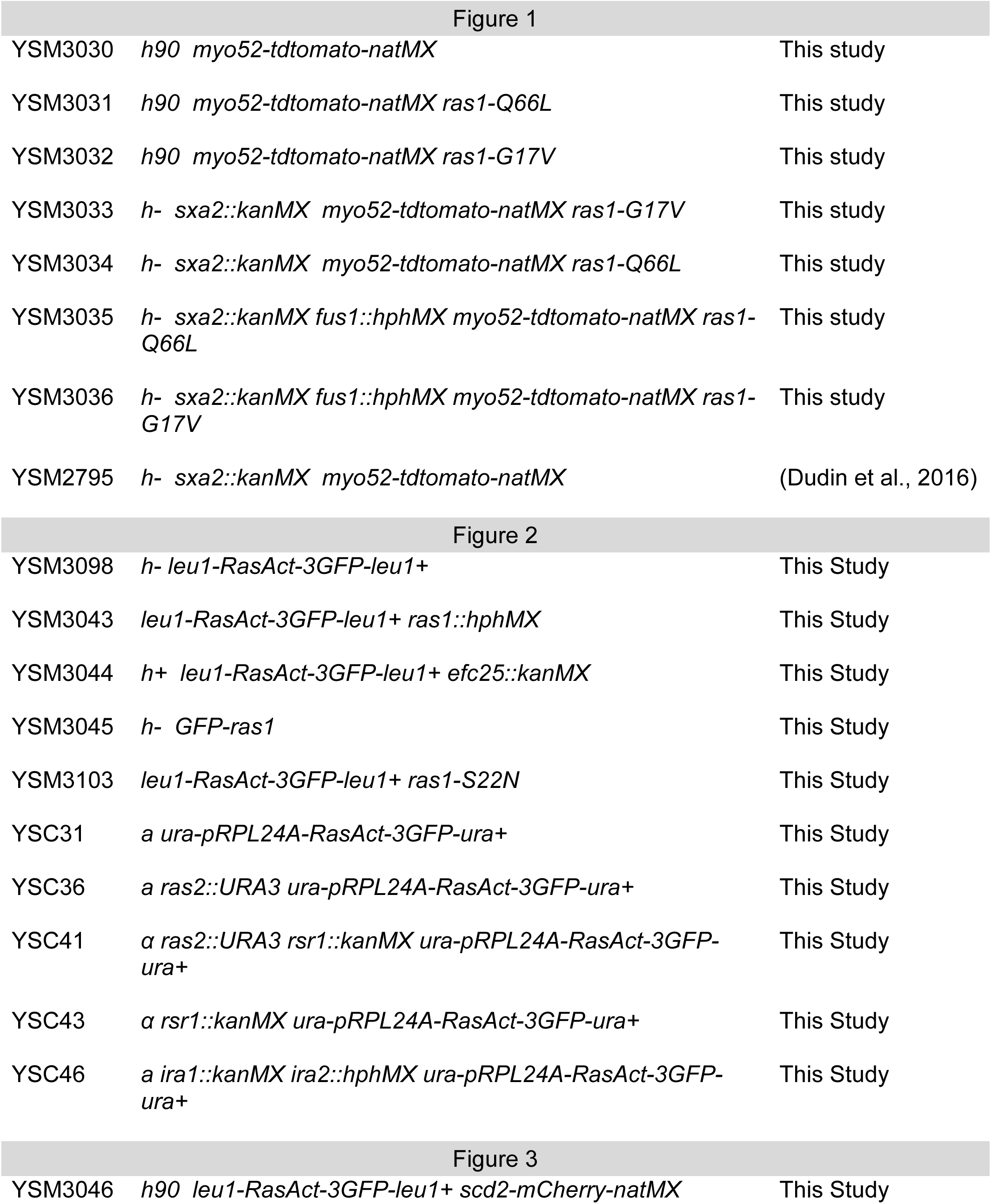
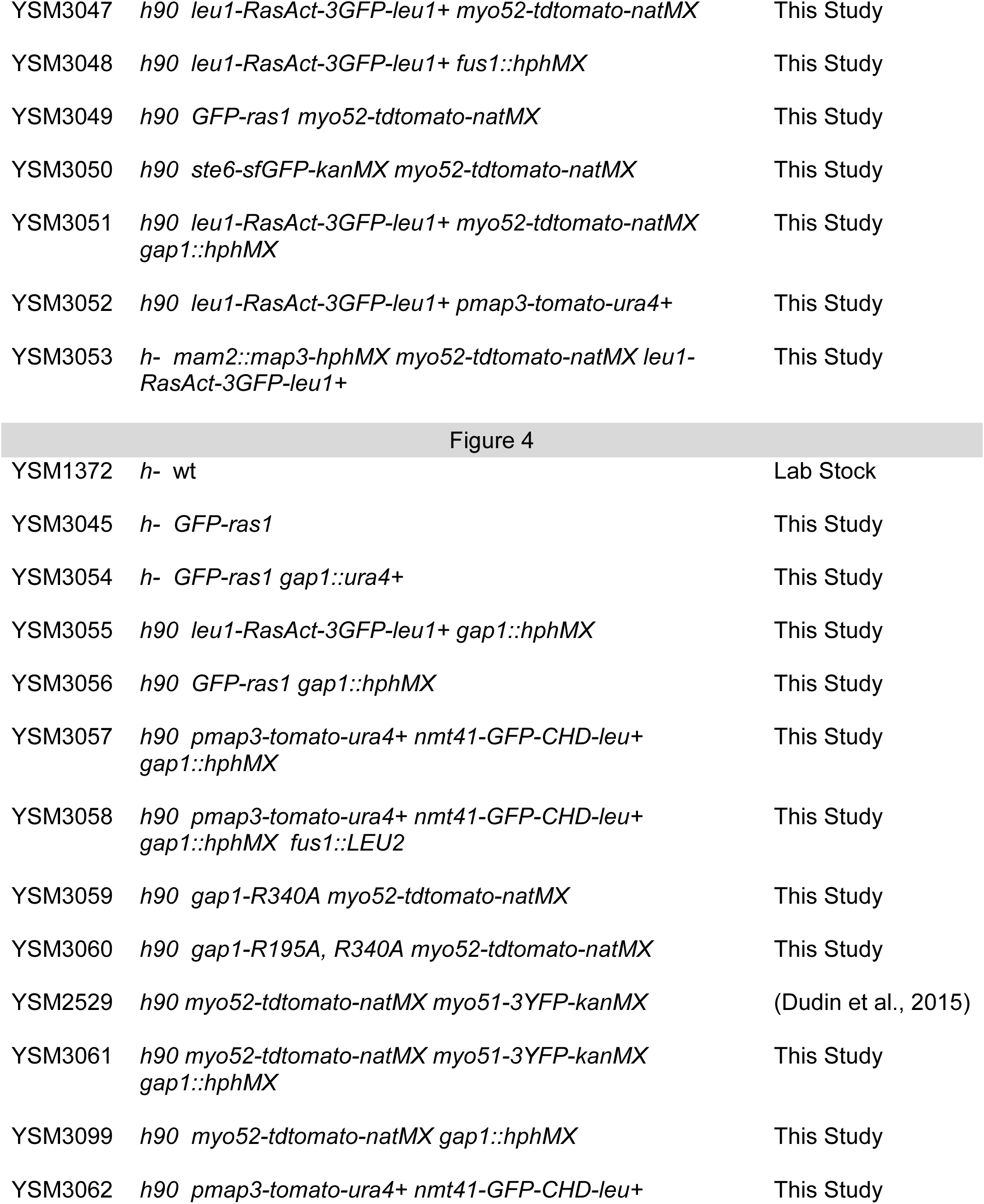
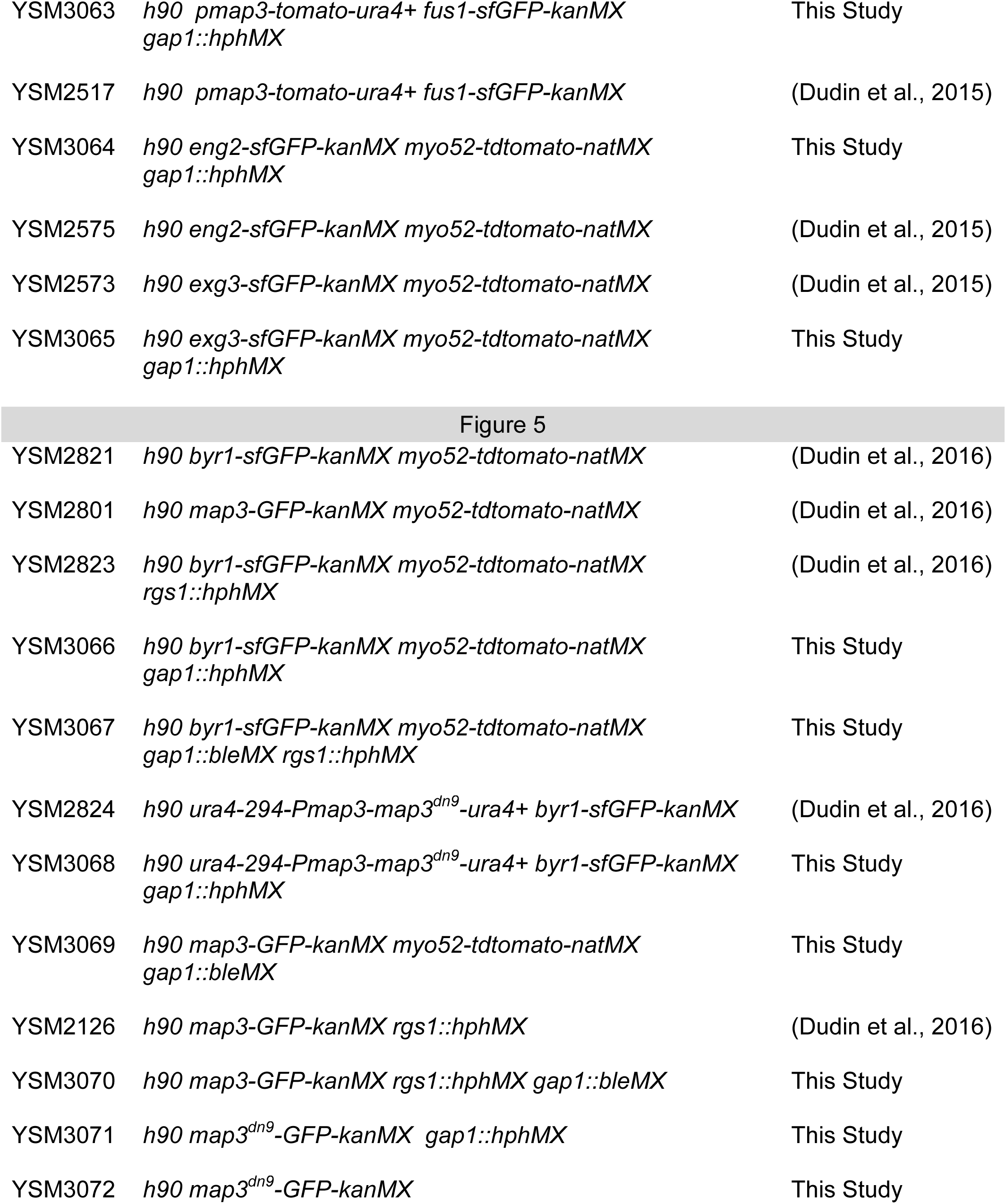
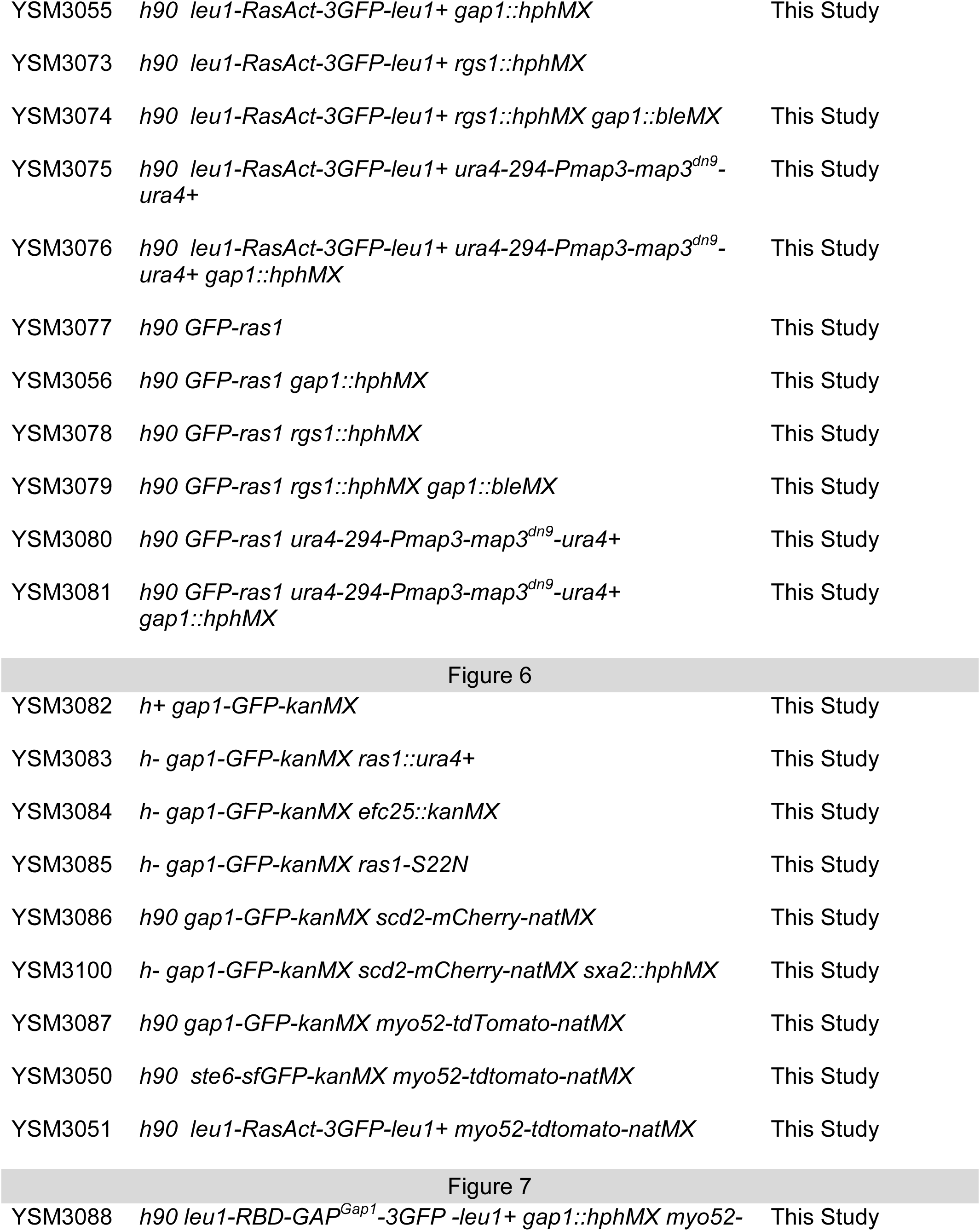
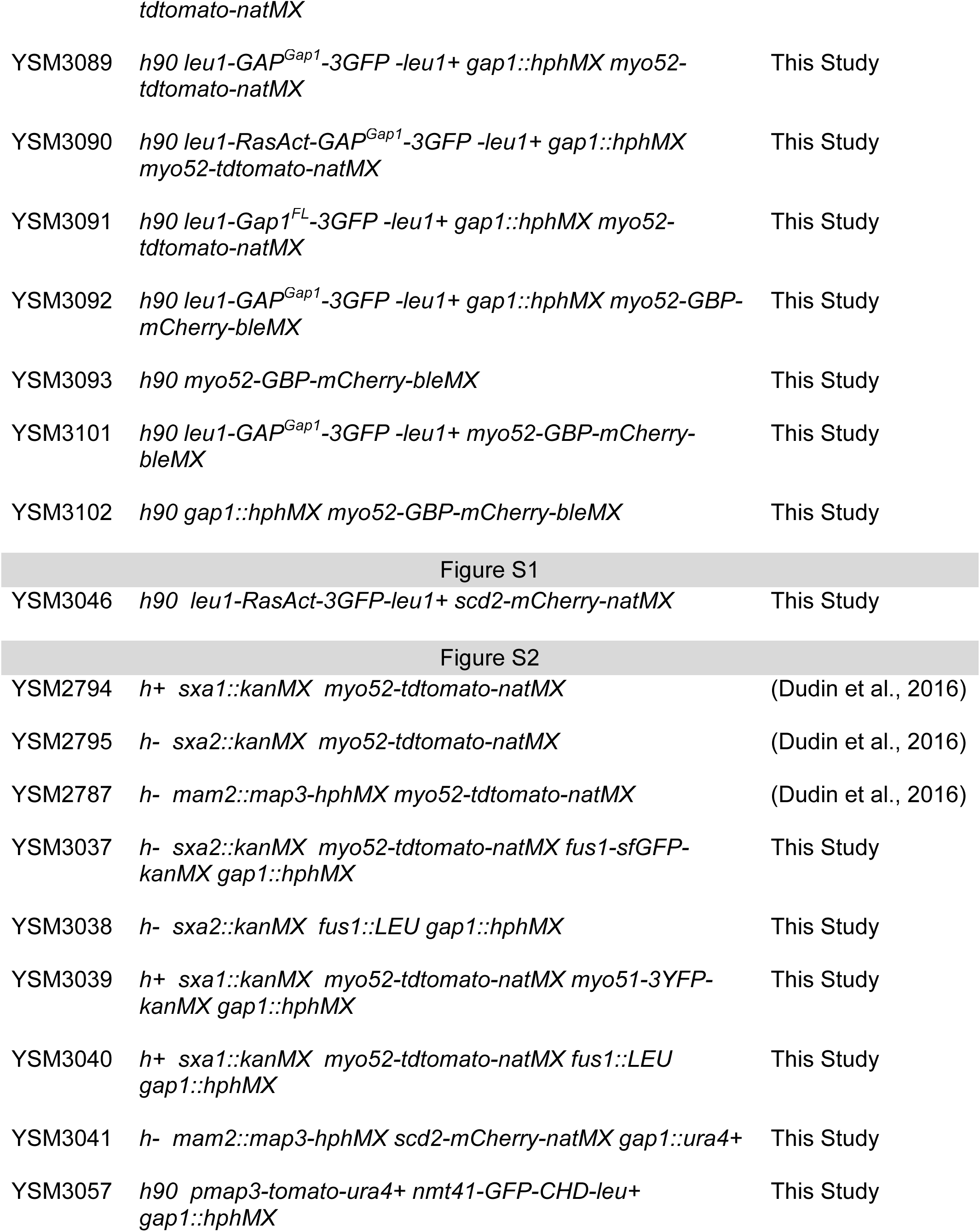
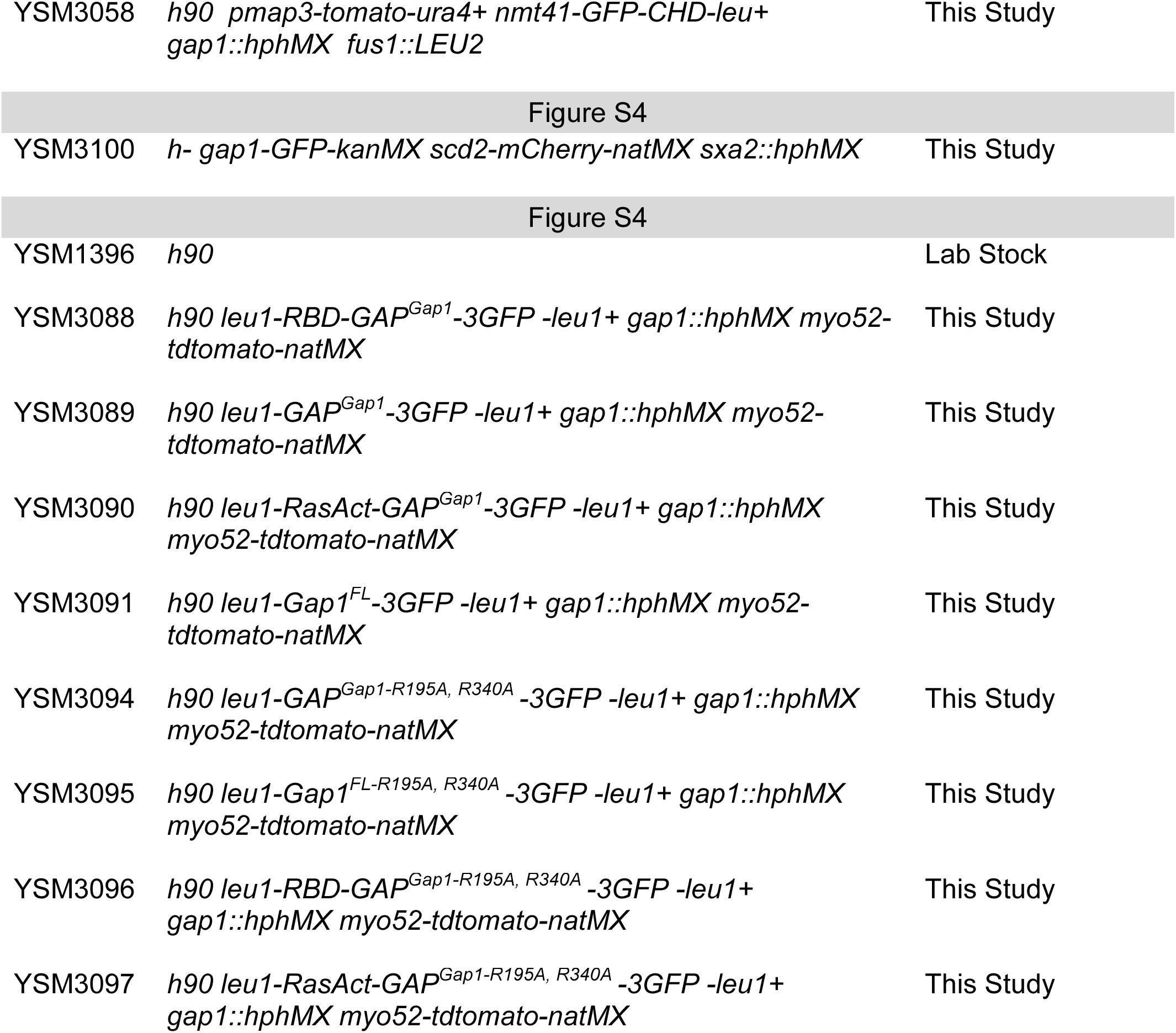
Strains used in this study.

